# Comparative transcriptomics identifies differences in the regulation of the floral transition between Arabidopsis and *Brassica rapa* cultivars

**DOI:** 10.1101/2020.08.26.266494

**Authors:** Alexander Calderwood, Jo Hepworth, Shannon Woodhouse, Lorelei Bilham, D. Marc Jones, Eleri Tudor, Mubarak Ali, Caroline Dean, Rachel Wells, Judith A. Irwin, Richard J. Morris

## Abstract

The timing of the floral transition affects reproduction and yield, however its regulation in crops remains poorly understood. Here, we use RNA-Seq to determine and compare gene expression dynamics through the floral transition in the model species *Arabidopsis thaliana* and the closely related crop *Brassica rapa*. A direct comparison of gene expression over time between species shows little similarity, which could lead to the inference that different gene regulatory networks are at play. However, these differences can be largely resolved by synchronisation, through curve registration, of gene expression profiles. We find that different registration functions are required for different genes, indicating that there is no common ‘developmental time’ to which Arabidopsis and *B. rapa* can be mapped through gene expression. Instead, the expression patterns of different genes progress at different rates. We find that co-regulated genes show similar changes in synchronisation between species, suggesting that similar gene regulatory sub-network structures may be active with different wiring between them. A detailed comparison of the regulation of the floral transition between Arabidopsis and *B. rapa*, and between two *B. rapa* accessions reveals different modes of regulation of the key floral integrator *SOC1*, and that the floral transition in the *B. rapa* accessions is triggered by different pathways, even when grown under the same environmental conditions. Our study adds to the mechanistic understanding of the regulatory network of flowering time in rapid cycling *B. rapa* under long days and highlights the importance of registration methods for the comparison of developmental gene expression data.

## Introduction

In nature, flowering time is a critical factor in determining a plant’s reproductive success (Ims, 1990). In agriculture, the control of flowering is important for determining yield, and must be optimised to fit within the constraints of the growing season. For rapid cycling oilseed *Brassica rapa*, the growing season may be determined by environmental constraints, such as the need to avoid potentially damaging climactic conditions (Canola Council of Canada, 2013), or by land management constraints, such as the requirement to fit within an established crop rotation. Specifically, in north-eastern Bangladesh, demand for short-duration oilseed varieties is driven by the need to fit within a “T. Aman rice – mustard – Boro rice” cropping pattern requiring extremely fast developing mustard varieties which can reach maturity in less than 80 days (Md *et al*., 2016; Miah & Mondal, 2017).

Much of our current understanding of the genetic regulation of the floral transition stems from studies on the model plant *Arabidopsis thaliana*. In Arabidopsis, the transition from vegetative to inflorescence development of the apex is regulated by the complex interaction of hundreds of genes across multiple tissues (Bernier & Périlleux, 2005; Pajoro *et al*., 2014; Bouché *et al*., 2016a; Périlleux *et al*., 2019). These interactions comprise a gene regulatory network (GRN) for flowering, which is commonly divided into a number of parallel exogenous and endogenous signalling pathways (photoperiod, ambient temperature, autonomous, vernalisation and aging (Simpson & Dean, 2002; Andrés & Coupland, 2012; Bouché *et al*., 2016b; Hyun *et al*., 2019). Signals from these different pathways are integrated at the apex to moderate timing of the floral transition, during which vegetative production of leaf primordia switches to production of floral primordia. This transition can be identified morphologically, and is also accompanied by changes in the expression of a number of well-characterised genes such as *FRUITFULL* (*FUL*), *SUPRESSOR OF OVEREXPRESSION OF CONSTANS* (*SOC1), LEAFY* (*LFY*) and *APETELA1* (*AP1*) (Klepikova *et al*., 2015).

In Arabidopsis, exogenous signals include photoperiod and temperature which are perceived primarily in the leaf. Under inductive environmental conditions, these signals culminate in the production of the mobile protein FLOWERING LOCUS T (FT). FT is able to move through the phloem to the apex where it activates flowering (Corbesier *et al*., 2007; Jaeger & Wigge, 2007). FT’s role as a signal of environmental conditions is similar in *B. rapa* (del Olmo *et al*., 2019). Conversely, in the perennial *Arabis alpina*, and *Arabidopsis thaliana* when grown under non-inductive conditions, shoots and branches can undergo the floral transition in the absence of *FT* expression, mediated by the independent endogenous aging pathway (Hyun *et al*., 2019).

*B. rapa* and Arabidopsis are both members of the Brassicaceae family, having diverged from their last common ancestor about 43 Mya (Beilstein *et al*., 2010). Given this relationship, it is likely that orthologues of the Arabidopsis genes play similar roles in *B. rapa*, and indeed *FLOWERING LOCUS C* (*FLC*), *FT* and *SOC1* orthologues have been identified as strong candidates underlying variation in flowering time in rapid cycling *B. rapa* (Lou *et al*., 2007; Franks *et al*., 2015; Zhang *et al*., 2015). However, differences in the expression dynamics of floral transition genes, both between Arabidopsis and *B. rapa* and between *B. rapa* accessions, remain largely uncharacterised, and the regulatory interactions controlling the floral transition in *B. rapa* remain poorly understood (Xiao *et al*., 2013; Blümel *et al*., 2015). It should be possible to use fundamental insights from Arabidopsis development to short-cut a similar understanding of development in related crop species. However, direct comparison between organisms is not trivial and it remains unclear to what extent their progression through development is comparable. Developmental differences may originate at any level from environmental perception onwards and cascade through the GRNs, making it difficult to distinguish original, causal differences from their consequences. Transcriptomics allows a broad survey of the behaviour of the regulatory system from which causal differences in developmental gene expression can be identified.

Here, we have generated extensive transcriptomic datasets for two oilseed *B. rapa* accessions. These comprise a time-course of gene expression in leaf and apex tissues beginning during vegetative growth and continuing through the floral transition until buds are visible on the plant. We compared gene expression between these varieties, and to publicly available rapid cycling Arabidopsis (Col-0) apical gene expression data (Klepikova *et al*., 2015).

R-o-18 is a commonly used laboratory accession, closely related to *B. rapa* oilseed crops grown in Pakistan (Rana *et al*., 2004). Sarisha-14 is a commercial cultivar developed at the Bangladesh Agriculture Research Institute (BARI) from local varieties. It develops extremely rapidly, reaching maturity in approximately 75 days, and is thus viable in a “rice-mustard-rice” cropping cycle (Md *et al*., 2016; Mia, 2017). Comparison to this unusual accession is carried out to identify commercially relevant GRN divergence in Sarisha-14 from more conventional rapid cycling oil type *B. rapa* accessions.

Transcriptome comparison between Arabidopsis and *B. rapa* by alignment of gene expression profiles using curve registration (Ramsay & Silverman, 2005; Leiboff & Hake, 2019) suggests that there is not one, but many different ‘developmental progressions’ of gene expression running at different speeds relative to each other. There is, therefore, no single common ‘developmental time’ in these closely related plants. In addition, we perform a detailed comparison of differences in the gene regulatory networks controlling flowering time. We find differences in the regulation of the apical expression of the transcription factor *SOC1* between Arabidopsis and *B. rapa*. Our data suggest an *FT* independent mechanism for extremely rapid flowering under long-day conditions in *B. rapa* in Sarisha-14, distinct from that present in rapid flowering R-o-18.

## Methods

### Plant growth conditions, sampling & gene expression quantification

*Brassica rapa* cv. Sarisha-14 (F_8_) and R-o-18 (double haploid) plants were sown in cereals mix (40 % medium grade peat, 40 % sterilised soil, 20 % horticultural grit, 1.3 kg m^−3^ PG mix 14-16-18 + Te base fertiliser, 1 kg m^−3^ Osmocote Mini 16-8-11 2 mg + Te 0.02 % B, wetting agent, 3 kg m^−3^ maglime, 300 g m^−3^ Exemptor). Material was grown in a Conviron MTPS 144 controlled environment room with Valoya NS1 LED lighting (250 μmol m^-2^ s^-1^) 18 °C day/ 15°C night, 70 % relative humidity with a 16 hr day. Sampling of Sarisha-14 and R-o-18 leaf and apex was performed 10 hrs into the day. Leaf (1st true leaf) and apex samples were taken over development during vegetative growth and the floral transition, continuing until floral buds were visible (developmental stage BBCH51, Meier *et al*., 2009). Each sample at each timepoint consists of pooled tissue dissected from leaf or apex of three different plants. Leaf and apex are taken from the same plants.

All dissections were performed on ice within the growth chamber and material harvested into LN_2_, prior to −70 °C storage. Samples were ground in LN_2_ to a fine powder before RNA extraction including optional DNase treatment was performed following the manufacturers standard protocol provided with the E.Z.N.A^®^ Plant RNA Kit (Omega Bio-tek Inc., http://omegabiotek.com/store/).

For *B. rapa* accessions, 150 bp paired-end RNA reads were generated at Novogene (Beijing). cDNA libraries were constructed using NEB next ultra-directional library kit (New England Biolabs Inc) and sequencing was performed using the Illumina HiSeq X platform. Publicly available gene expression data in *A. thaliana* Col-0 shoot apex from 7 days to 16 days after germination grown under similar 16 hr day conditions were downloaded from NCBI SRA, project ID PRJNA268115 (Klepikova *et al*., 2015). Gene expression quantification was carried out using HISAT v2.0.4 (Kim *et al*., 2015), & StringTie v1.2.2 (Pertea *et al*., 2015). Reads were aligned to either the *B. rapa* chiifu v3 reference genome (Zhang *et al*., 2018) (R-o-18, Sarisha-14), or the TAIR10 reference genome (Berardini *et al*., 2015) (Col-0). Gene expression level is reported in TMM normalised counts per million (Robinson *et al*., 2010).

### Comparison of gene expression states in biological samples

For comparison between Arabidopsis, and R-o-18, pairs of homologues with highly varied expression over development were identified. The criterion was that the correlation coefficient between expression of replicates at each timepoint, and mean expression over biological replicates at each timepoint was greater than 0.7 in both genotypes. This identifies genes for which mean expression over time changes by a large amount compared to variation between replicates at each timepoint. This selects 2346 *B. rapa*, and 1529 Arabidopsis homologues. The gene expression distance between timepoints for each pairwise comparison was calculated as the mean squared difference in gene expression between pairs of homologues. Gene expression scaling was carried out by subtracting mean expression over the time-course and dividing by the standard deviation in a gene-wise manner. Differential gene expression analysis between R-o-18 and Sarisha-14 vegetative apices was carried out using EdgeR (Robinson *et al*., 2010). t-SNE analysis (in which pairwise gene expression distances between samples were projected onto one dimension, whilst optimally retaining between sample distances) was used to compare development between R-o-18 and Sarisha-14, and was carried out using Rtsne (van der Maaten & Hinton, 2008; Krijthe, 2015).

### Registration of gene expression profiles over time

In order to register (align) gene expression profiles in Arabidopsis and *B. rapa*, Arabidopsis gene expression profiles over time were stretched and translated, using the least squares criterion to determine optimality. Specifically, gene expression levels were centred and scaled using the mean and standard deviation of the overlapping registered time points in each species. Stretch factors of 1x, 1.5x, and 2x, and translation factors between −4 and +4 days were considered. Stretching over only an arbitrary subsection of the observed timeseries was not considered, in order to minimise overfitting. After a candidate registration function was applied, gene expression was linearly imputed between the mean observed value at each timepoint in each species. For each gene, considered registrations were scored using the mean square difference between *B. rapa* observed timepoint, and the imputed Arabidopsis expression value over the overlapping timepoints. The best set of registration factors for each gene minimised this score and were carried forward to compare to a no-registration model. Bayesian model selection was used to compare the support for a no-registration model (in which expression over time for each gene is different between the two species) versus a registration model (in which expression profile differences can be resolved through the described registration procedure). For the overlapping timepoints identified after registration, cubic spline models with 6 parameters were fit; to expression in each species separately (2 × 6=12 parameters), or a single spline for gene expression in both species after the optimal “shift-and-stretch” registration transformation had been applied (2 + 6=8 parameters). The Bayesian Information Criterion (BIC) statistic was used to compare these models for each gene.

### Assortative mixing of registration parameter groups in gene network

Registration parameter groups were mapped to the AraNet v2 co-functional gene interaction network (Lee *et al*., 2015). The assortativity coefficient (a measure of the degree of mixing between genes with different optimal registration function parameters) was calculated by equation 7.80 of (Newman, 2010), and compared to 100,000 values calculated when the identified parameter groups were randomly allocated over the network, in order to estimate statistical significance.

### Network inference

The likelihood of regulatory links between genes were inferred from gene expression data following the Causal Structure Identification (CSI) algorithm (Penfold & Wild, 2011). The performance of this approach for data similar to ours was evaluated using synthetic gene expression data generated using networks of known structure, with varied experimental noise, correlation between candidate parents, generative GP hyperparameters. and numbers of observations (**Supporting Information Fig. S1**).

### Identification of pri-RNA homologues

Arabidopsis and *B. rapa* precursor-mRNA sequences, were downloaded from TAIR (https://www.arabidopsis.org/) and miRBase (Griffiths-Jones *et al*., 2006). Candidate pri-mRNA gene regions were identified in the Chiifu v3 reference sequence (Zhang *et al*., 2018) based on BLAST similarity (E-val < 1E-20), (**Supporting Information Table S1**). Stringtie v1.2.2 was used to reannotate the reference sequence using sequencing data from all Sarisha-14 and R-o-18 apex and leaf samples (**Supporting file S1**), and gene models overlapping the BLAST sites were considered to be candidate pri-RNA genes.

## Results

### Transcriptomes over development appear to be dissimilar between Arabidopsis and *B. rapa*

We compared gene expression across time, through the floral transition, in apical tissue of Arabidopsis ecotype Col-0 and *B. rapa* accession R-o-18. These closely related species move through a similar morphological sequence of developmental stages, so one might expect their transcriptomes to progress along a path of similar gene expression states. Under this assumption, we would expect to see that plants at similar morphological developmental stages exhibit similar transcriptomes (Leiboff & Hake, 2019).

To reduce noise and highlight differences and similarities in changes in developmental gene expression, we enriched the compared gene set for genes whose expression was found to change over the time-course relative to variability between biological replicates (see methods), resulting in comparison of 2346 *B. rapa* to 1529 Arabidopsis homologues.

Within each species, samples taken at similar times in general have more similar gene expression than samples taken at dissimilar times (**Fig. 1 a)**, indicating that our data is of a sufficient temporal resolution to detect developmental changes in transcriptome expression, and so identify similar developmental states. The exception to this is Col-0 day 11, which appears highly dissimilar to all other observed timepoints, and may represent an unusually short-lived gene expression state.

**Figure 1:**
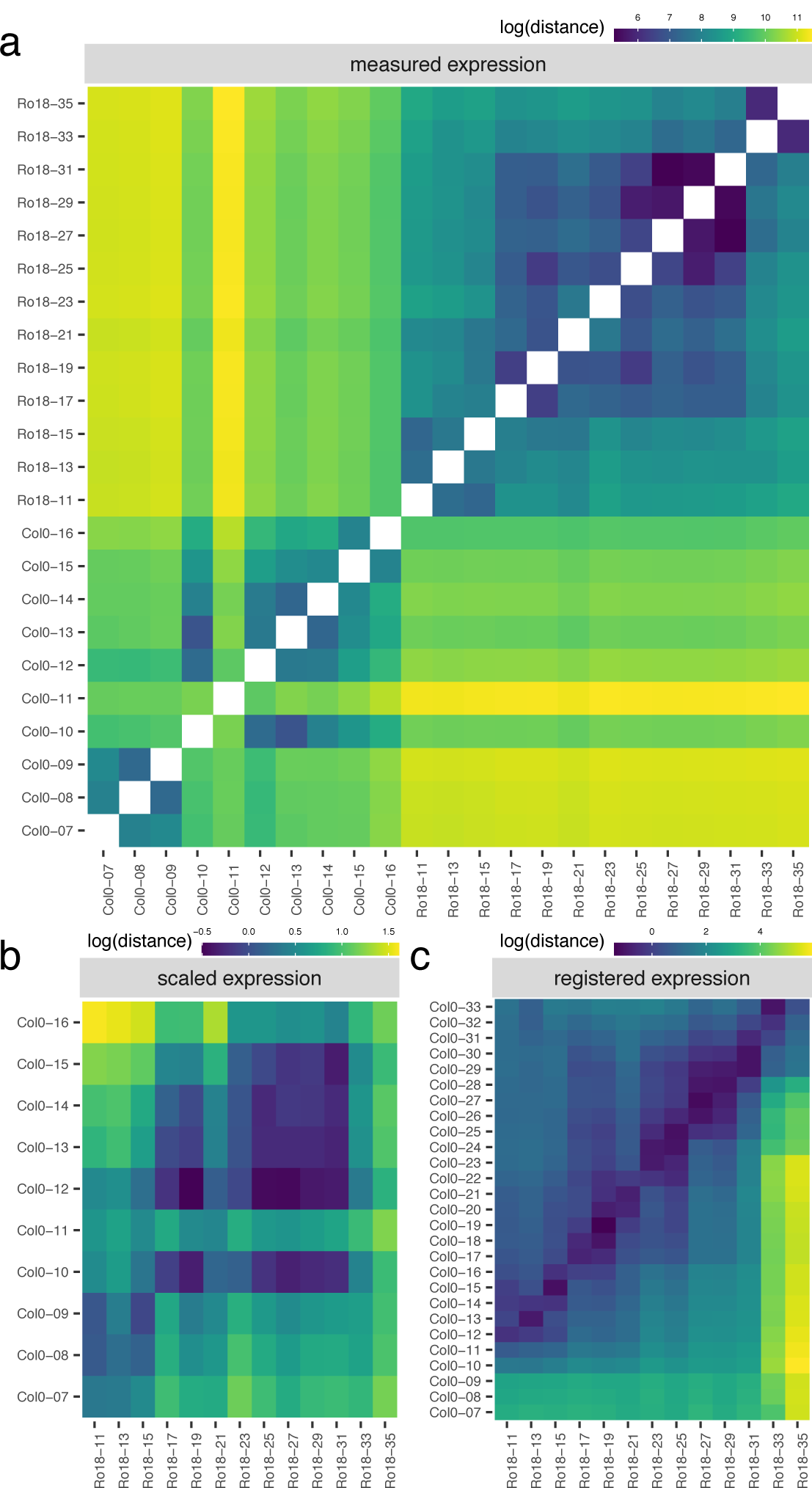
Gene expression states differ during development between Arabidopsis and *B. rapa* in the shoot apex. Heatmaps show the gene expression distance between samples taken from the apex of R-o-18 or Col-0 at varying days post germination. Gene expression distance between pairs of samples is calculated as the average squared difference in expression between homologous pairs of genes. **(a)** Measured gene expression counts are not similar between species over time. For comparisons made within each genotype (lower left, upper right quadrants), samples taken from points close in time (points near diagonal line) are more similar to each other than to samples taken from different times (points far from diagonal). Comparing between species (upper left and lower right quadrants), however, reveals no obvious structure. This suggests that species in similar morphological developmental states do not necessarily exhibit similar gene expression. **(b)** Scaled expression values are used to control for differences in magnitude. Note the change of axes from (1a) to compare only between species. In contrast to (1a), some diagonal structure is now apparent, reflecting some correspondence between expression at similar times in different species. **(c)** Bayesian model selection suggests that for many genes, differences between Col-0 and R-o-18 are more likely to stem from desynchronisation of the same expression patterns, rather than different expression patterns *per se* (see methods). The degree of desynchronisation differs between genes, and after this is accounted for, similar gene expression states between R-o-18 and Col-0 become apparent (block structure along the diagonal). This shows that there is a common progression through more gene states than just the blocks evident in (b).

To our surprise, however, no similarity can be seen in the progression of gene expression states between species (**Fig. 1 a**, upper left, lower right quadrants). The transcriptomes of the two species at points close in time do not appear to be more similar than the transcriptomes at more distant timepoints. This apparent lack of transcriptome similarity between organisms can be partly accounted for by differences in gene expression magnitude between organisms. After scaling gene expression in each organism (**Fig. 1 b**), later R-o-18 timepoints (from approximately 17 d) are more similar to later Col-0 timepoints (from approximately 10 d). However, the resolution at which similar stages can be seen is much less than within a species, as no developmental progression is obvious within these coarse “early” and “late” blocks.

Thus, despite their relatively close evolutionary relationship, this data suggests that gene expression dynamics during the floral transition may be quite different between Arabidopsis Col-0 and *B. rapa* R-o-18.

### Expression of key floral transition genes are similar, but differently synchronised in Arabidopsis and *B. rapa*

To check whether this apparent dissimilarity is due to confounding effects from genes whose expression is not involved in development, we examined the expression of key genes involved in regulation of the floral transition, and whose expression pattern is diagnostic for different developmental stages in Arabidopsis (Klepikova *et al*., 2015). In Arabidopsis, when SOC1 protein expression is induced in the shoot apex, SOC1 and AGL24 directly activate expression of *LEAFY* (*LFY*), a floral meristem identity gene. *AP1* is activated mainly by FT (expressed predominantly in the leaf, so not compared here), and is also necessary to establish and maintain flower meristem identity. When *LFY* and *AP1* are expressed, flower development occurs at the shoot apical meristem according to the ABC model, through the activation of genes such as *AP3* (Lee & Lee, 2010).

**Fig. 2a** shows that if only samples taken at the morphologically determined floral transition (vertical bar) are considered, expression of these key genes is similar in both species, suggesting that (as expected) these genes play a similar role in this transition in both species. However, when expression of these five genes are considered together over time, the timing of changes in each of their expression patterns are not the same in both organisms (at least under these experimental conditions), relative to the timing of changes in the other four genes. For example, in Col-0 *SOC1* expression starts to increase before *LFY*, and plateaus prior to the floral transition, whereas in R-o-18 both genes accumulate over the same period. *AGL24* expression peaks before the floral transition in Col-0, and after it in R-o-18. In Col-0 *AP1* expression increases rapidly during floral transition, whereas in R-o-18 it remains at a relatively low level until later in development. In Col-0 *AP3* expression increases rapidly one day after transition, whereas in R-o-18 there is no such increase within the first four days after the transition.

**Figure 2:**
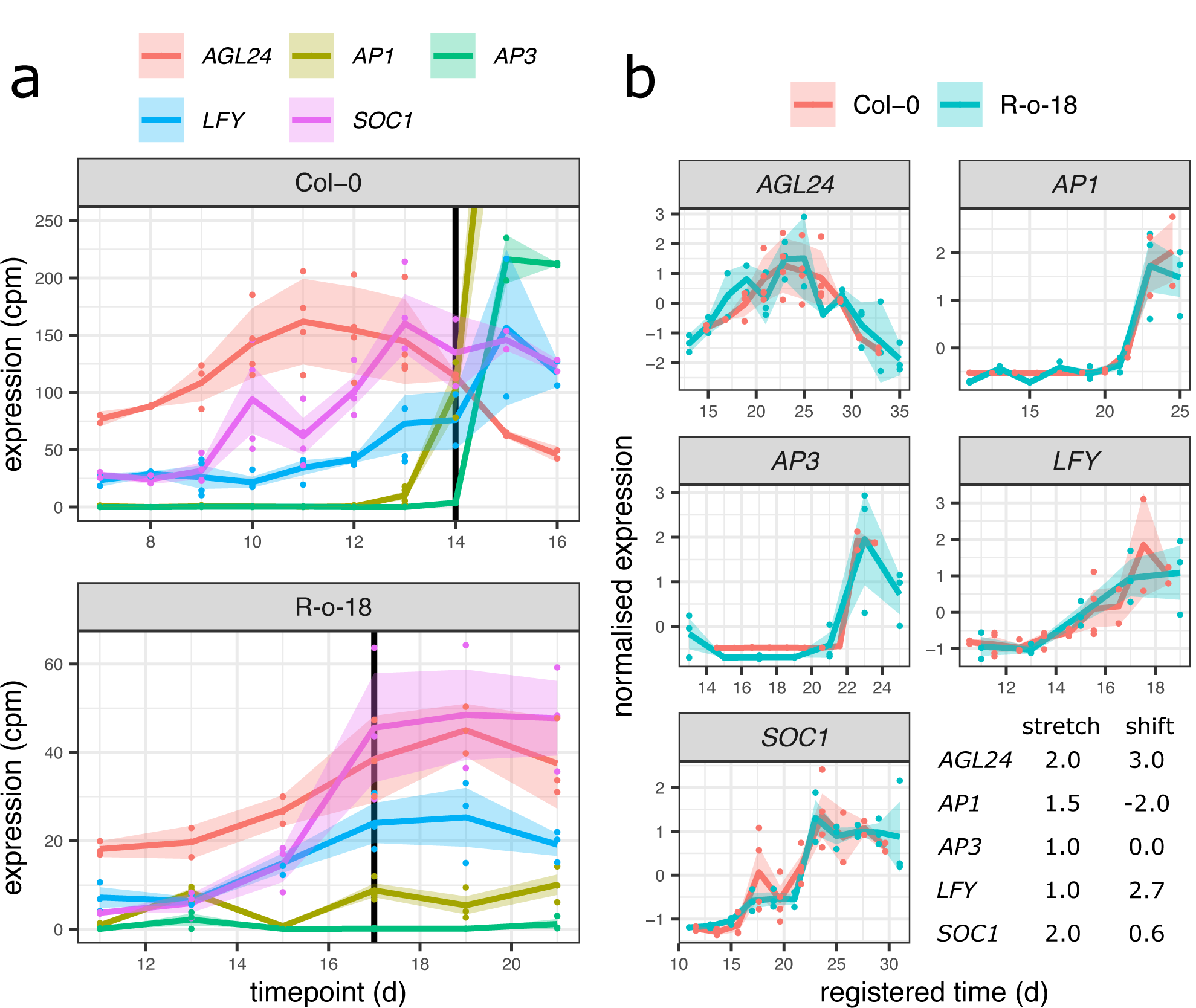
Key floral transition genes expression profiles are similar, but their timings are different between organisms. **(a)** Gene expression profile for five key floral transition genes in *A. thaliana* Col-0, and *B. rapa* R-O-18. Expression of paralogues in R-o-18 are summed. Morphologically identified floral transition time is identified by vertical line. The timings of gene expression changes relative to other genes, and the floral transition differ between R-O-18 and Col-0. **(b)** In spite of this, individual gene expression profiles are similar between these two organisms, as they superimpose after a registration transformation. The expression profiles of some genes are stretched out in R-o-18 relative to Arabidopsis (stretch), and also may be delayed, or brought forward relative to other genes (shift). The table shows the registration transformations applied to these genes; stretch indicates the stretch factor applied to Col-0 data, shift indicates the delay applied in days after this transformation.

To study and compare the expression dynamics of these genes in more detail, we employed curve registration (see methods). This method aims to synchronise functional data (here the gene expression over time of homologous pairs of genes in Arabidopsis and *B. rapa*) through the application of a suitable monotone transformation, translating and/or stretching gene expression profiles in an attempt to superimpose their dynamic behaviour.

**Fig. 2b** shows that following registration, the expression profiles of each pair of these exemplar genes can be superimposed and therefore have similar (though desynchronised) dynamics in Arabidopsis and *B. rapa*. This confirms our initial expectation that the expression of homologous genes might be similar between the two species. It shows that the differences in the expression profiles of these key gene pairs are differences in the relative timing, rather than in the nature or order of expression changes.

As can be seen in the table of optimal transformation function parameter estimates (within **Fig. 2b**), some differences in gene expression profiles between species are found to be explained by a shift (translation) in their expression over time (e.g. *LFY*), some are found to be explained by a stretch (e.g. *SOC1*), and some require a combination of these two factors. Also, the optimal amount of shifting and stretching differs between genes. Differences in the optimal registration function parameters of different genes highlight that the expression patterns of these individual genes are not desynchronised by the same amount between species.

Different delays in the timing of each gene’s expression means that (at least for this small set of genes) the expression of the combined set of genes is, in general dissimilar to any single timepoint in the other species. This is the case even though the expression patterns over time of the individual genes within this set are highly similar between species. When a larger set of genes (e.g. the whole transcriptome) is compared at single time points, these differences are likely to become more pronounced.

### Differences in the relative timing of gene expression changes between *B. rapa* and Arabidopsis are common

In order to evaluate the extent to which desynchronised expression changes might explain the apparent difference in transcriptomic gene expression, we applied the same registration procedure to the full set of genes which were found to vary in expression over the time-course. We found that for 1465 of the 2346 considered *B. rapa* genes, the Bayesian Information Criterion (BIC) favours a model that considers gene expression in *B. rapa* and *A. thaliana* to be the same (after registration) over a model in which they are considered to have different gene expression patterns. Permutation testing, in which genes in one organism are randomly allocated a comparison gene in the other, suggests that this is a significantly large number of genes to be identified (p < 2e-23, **Supporting Information Fig. S2**), and therefore not merely an artefact of overfitting during registration.

This analysis supports the conclusions drawn from the close examination of the few key floral genes and identifies differences in synchronisation as a general phenomenon. Thus, for many genes, the difference between R-o-18 and Arabidopsis is a delay in the gene’s expression pattern, rather than a more complicated difference in their expression dynamics.

When these differences in timing are accounted for (through registration), there is a further reduction in the distance between nearby timepoints, and increase in the distance between dissimilar timepoints (**Fig. 1c**). The heatmap shows a common progression from early to late gene expression states in both species. This indicates that gene expression over time is much more similar between these organisms than could be concluded through a naïve comparison of their gene expression profiles over time without registration. It partially resolves the apparent paradox that *B. rapa* and Arabidopsis are related organisms with highly similar morphological development, but which apparently exhibit dramatically diverged gene expression patterns over development even when grown under similar environmental conditions.

As in the floral gene example, different optimal registration transformation parameters are identified for different genes (**Supporting Information Fig. S3**). Contrary to our initial hypothesis, it is therefore not the case that there is a single progression through transcriptomic states at different rates in *B. rapa* and Arabidopsis which could be aligned between them. Rather, there are a number of progressions bound together within each organism. These are each moved through at different rates, and only when they are synchronised through different registration functions, can we see how similar they are in both species. Thus, we find that there is not, in general, an equivalent developmental stage at the transcriptome level, and therefore no way to map both Arabidopsis and *B. rapa* to a single common developmental time in terms of overall gene expression.

### Differently synchronised groups of genes correspond to biologically functional groups, and position in gene regulatory network

In order to identify whether known biological GRN features correspond to these differently synchronised progressions, we examined groups of genes with the same optimal registration parameters, and which therefore exhibit synchronised expression differences between the *B. rapa* and Arabidopsis time-courses. Interestingly, groups of genes with the same optimal registration parameters are enriched in the same gene ontology terms, suggesting they may be involved in similar functions and processes (**Supporting Information Table S2**). Furthermore, when superimposed over an Arabidopsis gene-gene interaction network (Lee *et al*., 2015), genes in the same registration parameter group are more frequently linked to each other than to genes in a different parameter group (p<6e-5). Together these findings indicate that synchronised gene groups are associated with functional modules within the gene regulatory network.

That many genes have a similar expression patterns in both organisms, with co-functional genes co-synchronised within each organism, indicates that in general regulation of genes are highly similar in both Arabidopsis, and *B. rapa*. It suggests that under these environmental conditions, the GRN in *B. rapa* can be usefully understood as modules of genes with highly similar regulatory relationships as in Arabidopsis (resulting in their co-synchronisation), and that relatively few differences in gene-gene regulatory relationships, or environmental inputs leads to desynchronisation between these modules, and differences in expression.

### Regulation of *SOC1* differs between Arabidopsis and R-o-18

To further characterise an example of a gene regulatory difference between Col-0 and R-o-18, we focus on the regulation of *SOC1* in the apical flowering time network. This transcription factor is involved in the regulation of the upstream stages of the floral transition, and (as shown in the **Fig. 2)** its expression pattern is stretched by a factor of two in *B. rapa* relative to Arabidopsis, meaning that it comes on later, relative to other genes, and is slower to progress through its expression changes. Therefore, any differences in the regulation of SOC1 which explain this delayed expression are promising candidates to be involved in the delayed floral development in *B. rapa* relative to Arabidopsis.

To investigate potential *SOC1* regulatory changes we derived statistical models for the GRN from the data using the Causal Structure Inference (CSI) algorithm (Penfold & Wild, 2011). Comparison of the probability of candidate gene-to-*SOC1* regulatory links based on gene expression profiles suggests that among the largest differences in the regulation of *SOC1* between Arabidopsis and R-o-18 are changes in the response to *FLC* and *FUL* expression (**Supporting Information Fig. S4**). **Fig. 3** shows that although in Arabidopsis expression of *SOC1* is consistent with regulation via repression by *FLC* and activation by *FUL* as proposed by Balanza et al. (Balanzà *et al*., 2014), in R-o-18 none of the copies of *FLC* strongly associate with *SOC1*. Instead, *SOC1* expression is strongly associated with expression of the two *FUL* paralogues located on chromosome A02 and A03 (BRAA02G042750.3C, BRAA03G043880.3C). To understand the reason for the missing inferred regulatory links between *FLC* and *SOC1*, we considered the expression of *FLC* in more detail. Of the four paralogues of *FLC* identified in *B. rapa, BraFLC*.*A02* (BraA02g003340.3C) and *BraFLC*.*A10* (BraA10g027720.3C) have previously been reported to be non-functional in R-o-18, (Yuan *et al*., 2009; Wu *et al*., 2012; Schiessl *et al*., 2017). Our data shows *BraFLC*.*A03b* (BraA03g015950.3C) is also likely to be non-functional as there is a premature stop codon in exon 2, and it is expressed at a similar level to the other non-functional copies (**Supporting Information Fig. S5**).

**Figure 3:**
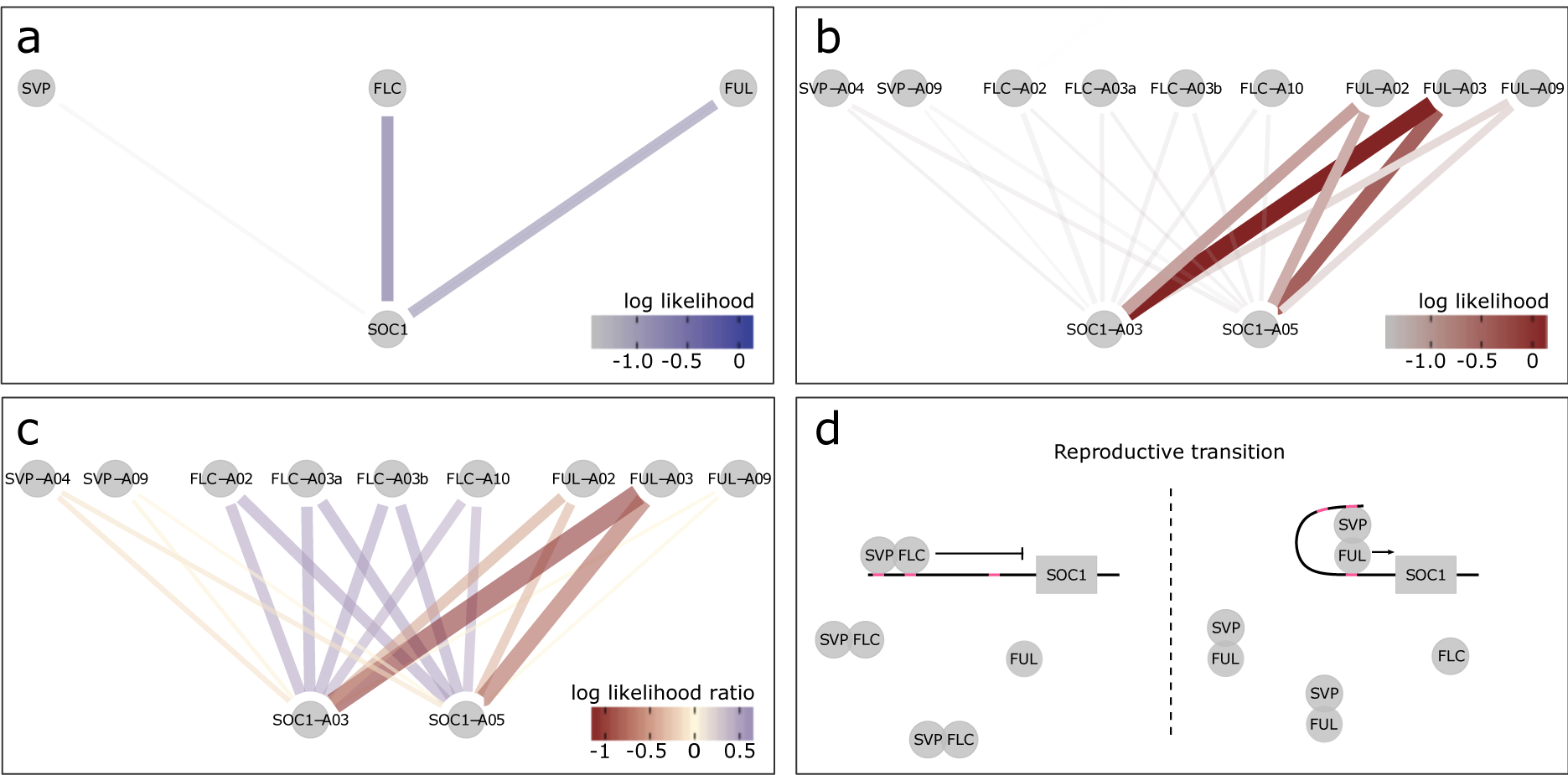
*SOC1* is differentially regulated between *B. rapa* R-o-18 and Arabidopsis Col-0. CSI inferred gene regulatory networks between *SVP, FLC, FUL* & *SOC1* in **(a)** Arabidopsis, **(b)** R-o-18. The likelihood of the observed gene expression data given an assumed regulatory link between each pair of genes is plotted. In the absence of prior information, this is proportional to the probability of a regulatory link between the gene pair given the observed gene expression data. **(c)** the difference between log likelihood in Col-0 and R-o-18. Numbers after gene abbreviation indicates chromosome number of the orthologue. **(d)** proposed mechanistic model for the role of FUL during the floral transition in Arabidopsis, modified from Balanzà *et al*. (2014), in which FUL and FLC compete to dimerise with SVP. In Arabidopsis, the CSI method infers that regulation of *SOC1* is via a balance of *FLC* and *FUL* expression, consistent with this model. Conversely in R-o-18, association *SOC1* is primarily between *SOC1*, and the A2 and A3 copies of *FUL* suggesting that changes in the expression level of FLC are not relevant to controlling the upregulation of *SOC1*.

*BraFLC*.*A03a* (BraA03g004170.3C) appears to be functional, is expressed at a higher level, and does not encode a premature stop codon. In Arabidopsis, apical *FLC* expression declines prior to *SOC1* upregulation, but in R-o-18 and Sarisha-14, *BraFLC*.*A03a* expression declines only after *SOC1* is upregulated (**Supporting Information Fig. S6**). This suggests a model such that in rapid cycling *B. rapa*, unlike rapid cycling Arabidopsis, the transition from vegetative to inflorescence meristem occurs prior to a decrease in expression of the floral repressor *FLC* in the apex. Consequently, the *SOC1* expression profile over development is delayed in R-o-18 relative to other flowering genes.

### The rates of development differ between leaf and apex in *B. rapa*

To evaluate whether comparison of transcriptomic timeseries could identify variation in GRNs underlying phenotypic variation between accessions, we compared gene expression in the leaf and apex of R-o-18 and Sarisha-14 *B. rapa* varieties. R-o-18 is a well-studied rapid yellow sarson oil type, Sarisha-14 is a commercially relevant rapeseed mustard, which develops extremely rapidly, undergoing floral transition 10 days after germination, 7 days earlier than R-o-18 (**supporting information Fig. S7**).

We computed the similarity in gene expression between different timepoints in R-o-18 and Sarisha-14 in leaf (**Fig. 4 a**), and apex (**Fig. 4 b**) tissues. This suggests that in the leaf, development overall proceeds at the same rate, as the most similar samples between accessions are at roughly equivalent timepoints. This is not the case in the apex, where there again appears to be a similar developmental trajectory in terms of gene expression, but progression along this path is faster in Sarisha-14 than in R-o-18. This desynchronisation of developmental progression suggests that differences in the rate of development between these accessions likely occur at the shoot apex, rather than the leaf, and implies that differences might exist in leaf to apex signalling of the floral transition between these accessions.

**Figure 4:**
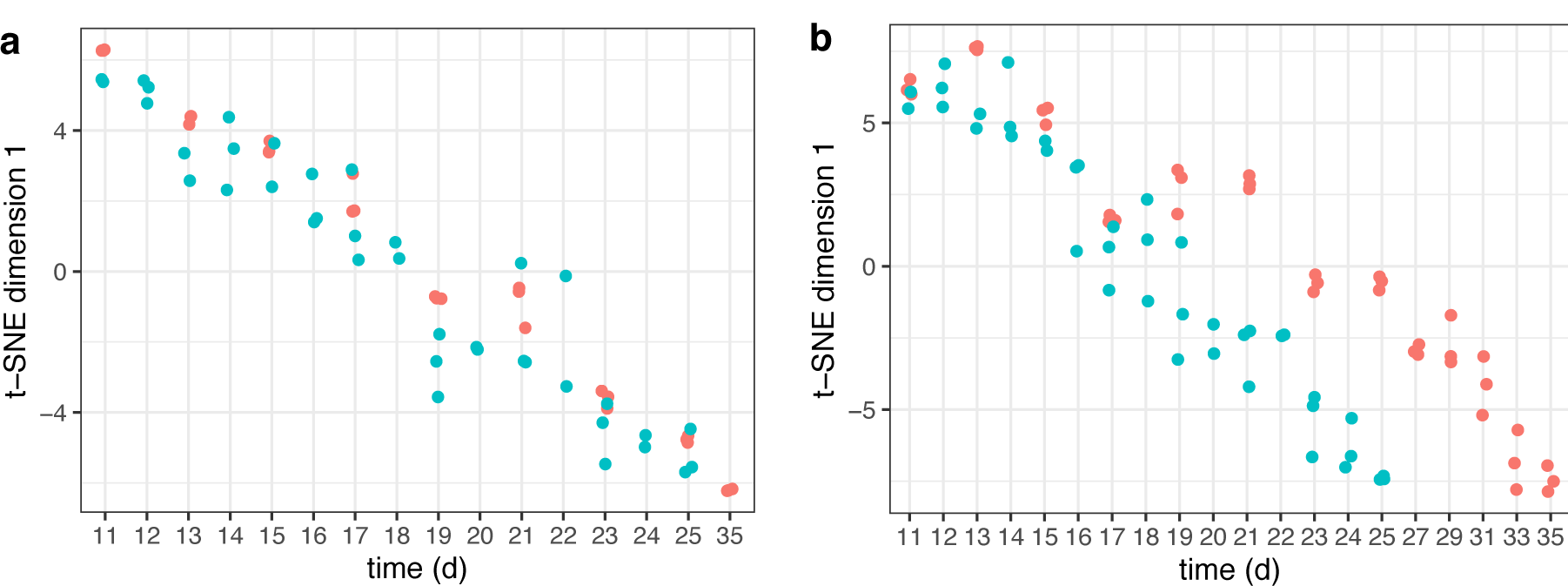
Developmental rates differ between Sarisha-14 and R-o-18 in the apex. Plots of time (days) against t-SNE estimated projection of gene expression to one dimension. This is an estimate of the optimal projection of the gene expression data whilst maintaining the correct distances between samples. Samples nearer to each other on the y-axis in each plot have more similar gene expression. Samples taken from **(a)** leaf and **(b**) apex, in R-o-18 (red), and Sarisha-14 (blue). In leaf, development of gene expression profiles over time appears to occur at approximately the same rate between accessions, such that the most similar samples are taken at the same time. In apex, development appears to occur faster in Sarisha-14 than R-o-18. Genes were filtered to only include genes which variation over time explained > 50% of variance in gene expression in both accessions. In apex, 3,097 genes were used. In leaf 10,035 genes were used.

### Rapid floral transition in Sarisha-14 is not due to an early *FT* signal, but to increased apical sensitivity

In Arabidopsis, environmental triggers of flowering are perceived predominantly in the leaf and result in the production of FT protein, which moves to the apex as a component of the florigen signal (Corbesier *et al*., 2007; Jaeger & Wigge, 2007). This then causes upregulation of flowering genes in the apex, such as *FUL*, and *SOC1* (Abe *et al*., 2005; Yoo *et al*., 2005; McClung *et al*., 2016). In *B. rapa, BraFT*.*A02* (BRAA02G016700.3C) which has previously been shown to be the main FT-like gene regulating the floral transition in R-o-18 (del Olmo *et al*., 2019), is the copy with the highest expression in both Sarisha-14 and R-o-18. In contrast, the *BraFT*.*A07* paralogue (BRAA07G031650.3C) contains a transposon insertion in R-o-18, which is predicted to generate a loss of function allele (Zhang *et al*., 2015) and is not detectibly expressed in either accession in our data. This suggests that it is not functional in either accession, and so is not considered here.

Meristems are floral seven days earlier in Sarisha-14 than R-o-18 (**Supporting Information Fig. S7**). However, registration indicates that *FT* expression in the leaf is only approximately 2 days ahead in Sarisha-14 compared to R-o-18. We also find that CSI inferred evidence for relationships between gene expression profiles in the leaf and the floral integrator genes *FUL* and *SOC1* in the apex is weaker in Sarisha-14 than R-o-18 (**Supporting Information Fig. S8**). In particular, less evidence for a relationship between *FT* expression in the leaf, and changes in apical gene expression were found in Sarisha-14 than R-o-18. Manual inspection of the expression pattern indicates that *FT* is not expressed sufficiently early in Sarisha-14 leaf relative to R-o-18 to account for the difference in the timing of the floral transition (**Fig 5**). It is not detectibly expressed prior to the floral transition, and at the time of floral transition, expression of *FT* in the leaf is lower in Sarisha-14 than R-o-18 (p=0.0272). Therefore, a given *FT* expression level in the leaf appears to result in a stronger induction of flowering response in Sarisha-14 than R-o-18. This could be achieved by: 1) increased potency of the FT signalling molecule; 2) increased conductance of the signal to the apex; 3) increased sensitivity of the apex to a signal; or 4) because floral transition occurs independently of *FT* in Sarisha-14. These can be considered as differences in signalling strength (models 1 & 2), and differences in signal perception at the apex (models 3 & 4). We find that gene expression at the apex is consistent with the third or fourth models.

**Figure 5:**
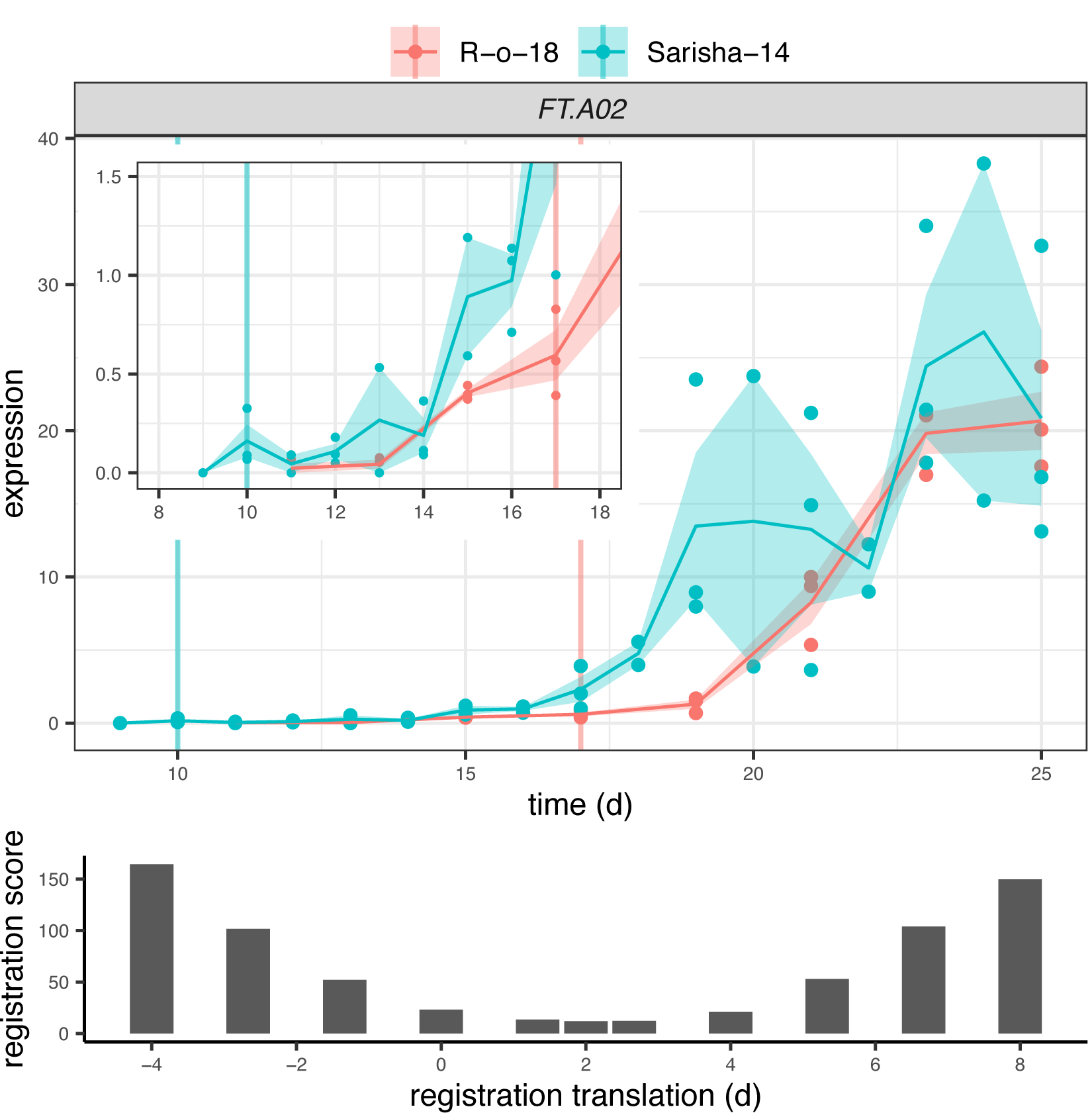
*FT* expression in Sarisha-14 leaf is not sufficiently early relative to R-o-18 to account for the difference in timing of floral transition. Gene expression of *BraFT* in R-o-18 and Sarisha-14 over development, inset graph shows expression before day 18. Vertical lines indicate the first timepoint with floral meristems identified in each accession. Registration indicates that expression of *FT* in the leaf is approximately 2d advanced in Sarisha-14 relative to R-o-18. This is not sufficient to account for the 7d difference in timing of the floral transition. Upon examination of the expression profiles, *FT* expression in the R-o-18 leaf increases between 13d and 15d, prior to floral transition at 17d. *FT* expression is not detectible in Sarisha-14 prior to the floral transition at 10d. Expression of *FT* in the Sarisha-14 leaf at floral transition is lower than in R-o-18 (17d). This shows that Sarisha-14 undergoes floral transition at the apex coincident with lower *FT* expression in the leaf than in R-o-18. It is not clear from this data whether *FT* is expressed in Sarisha-14 below the experimentally detectible limit prior to the floral transition. It is therefore unclear from this data whether the transition occurs in response to a reduced leaf FT signal, or even in its absence in Sarisha-14 grown under long-day conditions.

To identify any differences in apical gene expression which might cause increased sensitivity to an *FT* signal, or flowering in its absence, we compared apical gene expression in the last vegetative Sarisha-14 sample (9d post-germination), and the nearest vegetative R-o-18 timepoint (11d post germination). Both of these samples are prior to *FT* expression in the leaf, and so before differences in signal strength could affect behaviour. We found that 11,914 of 36,935 expressed genes which are differentially expressed (q<0.05), suggesting broad differences in gene expression. Among these genes, enriched representation of gene ontology terms “positive regulation of development, heterochronic” (q=0.017 FDR),” shoot system morphogenesis” (q=2.3e-4 FDR), and “phyllome development” (q=7.7e-4 FDR) indicate that developmental gene expression programs differ in the apex between these samples before they could be caused by *FT* signal strength differences.

We next investigated whether differences in other signalling pathways in the apex could account for the apparently different *FT* signal sensitivity. In Arabidopsis, the floral transition is controlled by multiple interacting pathways that are sensitive to environmental cues, as well as developmental age, which is controlled by a complex interaction between phytohormone signalling, sugar status, and the activity of microRNAs miR156 and miR172, and prevents premature flowering in juvenile plants. Signals from these different pathways are perceived and integrated at the shoot apex (**Fig 6**). We identified differently expressed genes in the miR156-SPL and AP2-like regulatory modules, but not in expression of *FLC, SVP*, or *FD*. In particular we note that that expression of miR156, miR172, and *SCHLAFMÜTZE* (*SMZ*), (the only AP2-like gene which is found to vary in transcriptional expression in response to perturbed miR156-miR172 expression (Yu *et al*., 2012)), are similar in Sarisha-14, and R-o-18 immediately prior to the floral transition in both accessions (**Fig 7**), though these events occur one week apart in time from germination.

**Figure 6:**
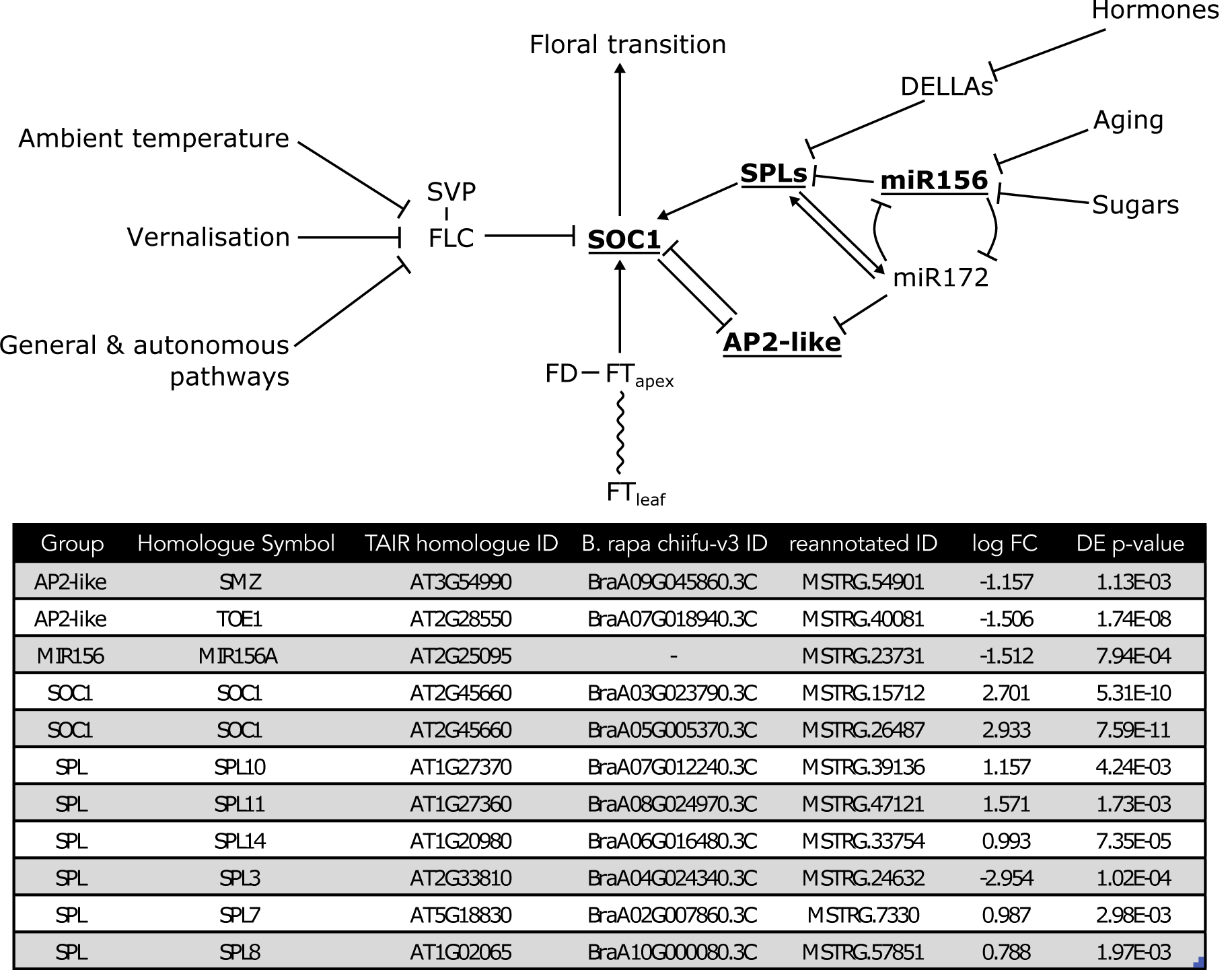
Differential expression at the convergence of multiple signalling pathways in regulation of the floral transition. Modified from the Flowering Interactive Database website (Bouché *et al*., 2016b), elements which were found to be differently expressed in the apex in pre-floral Sarisha-14 (day 9), and the nearest equivalent R-o-18 sample (day 11) are highlighted in bold and underlined. The table gives details of differently expressed gene identities, and log-fold change in Sarisha-14 relative to R-o-18. Differential expression of *SOC1* is coincident with differential expression of *SPLs* and AP2-like genes, rather than *FLC, FT, SVP*, or *FD*, implicating the endogenous Aging, Hormone, or Sugar signalling pathways in priming the early floral transition of Sarisha-14. Phytohormone signalling is integrated through the regulation of DELLA proteins. The activity of DELLA proteins is regulated post-translationally by GA, ABA, auxin, and ethylene either directly or indirectly (Fu & Harberd, 2003; Achard *et al*., 2006; Lorrai *et al*., 2018). Activity of SPLs are regulated by DELLA proteins (Conti, 2017). miR156 and miR172 are master regulators of the transition from the juvenile to adult phase of vegetative development (Wu & Poethig, 2006). During development initially high levels of mature miR156 and low levels of miR172 transition to low levels of miR156 and high levels of miR172, contributing to the juvenile to adult transition (Wu & Poethig, 2006; Hong & Jackson, 2015). miR156 primarily regulates SPLs via translational regulation (He *et al*., 2018). SOC1 is regulated by AP2-like transcription factors, and SPLs (Yant *et al*., 2010). AP2-like genes are regulated by the aging pathway, via largely via translational repression by miR172, though expression of the AP2-like gene *SMZ* has been found to depend on miR172 (Aukerman & Sakai, 2003; Chen, 2004; Yu *et al*., 2012).

**Figure 7:**
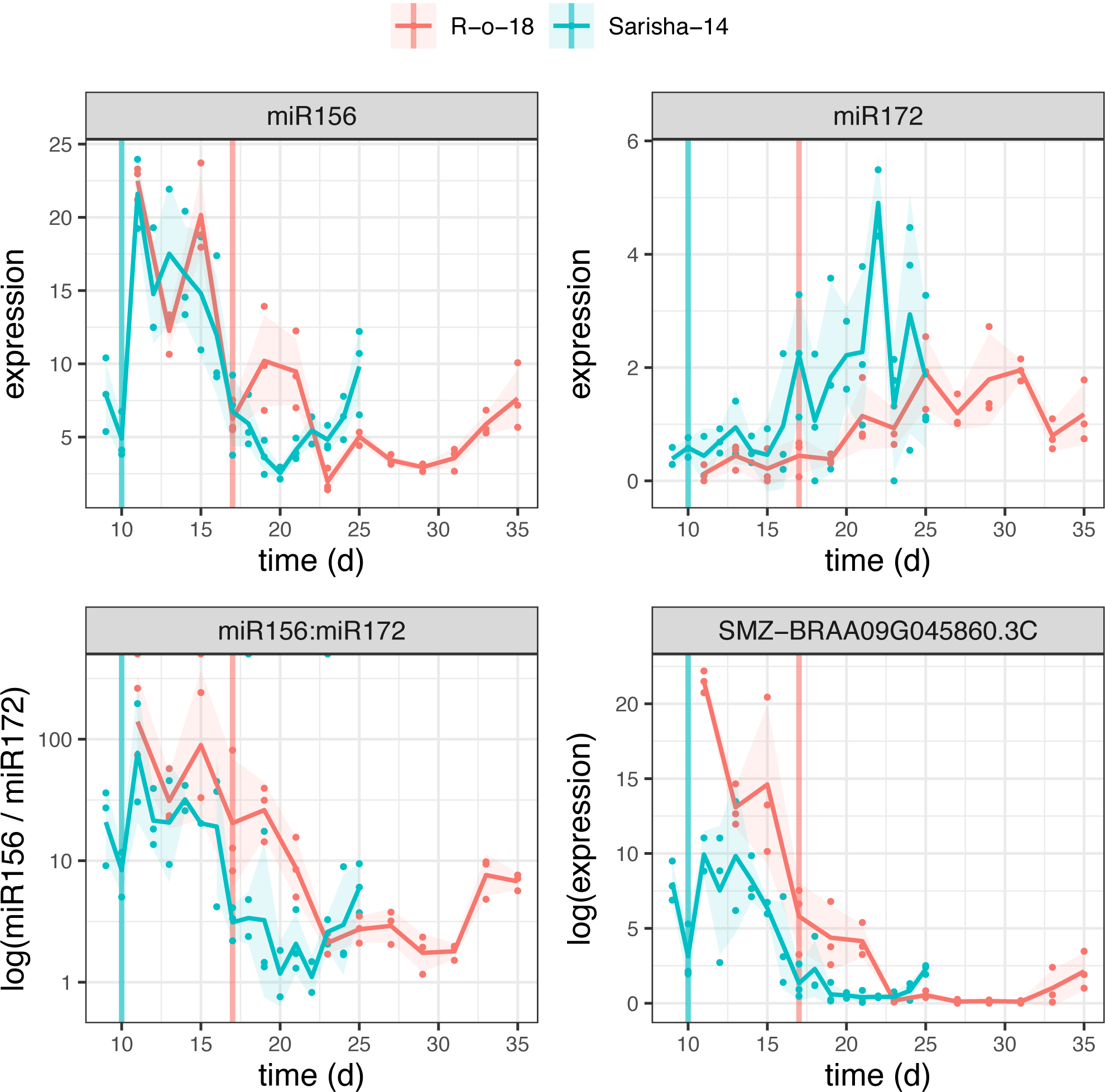
The aging pathway proceeds more rapidly in Sarisha-14 than R-o-18. Pri-miRNA gene models were identified as described in Methods. The ratio of miR156 to miR172 precursor RNA is lower in Sarisha-14 than R-o-18 at equivalent timepoints. This is achieved primarily though reduced expression of pri-miR156, though pri-miR172 is also expressed at a slightly higher level in Sarisha-14 than R-o-18. *SMZ* is transcriptionally regulated by miR172 (Yu *et al*., 2012), and so its lower expression level in Sarisha-14 suggests that miR172 activity as well as precursor levels are also greater in Sarisha-14. Mean and 95% CIs are shown.

These findings suggest that the early floral transition in Sarisha-14 is caused primarily by differences to R-o-18 in *FT* signal sensitivity at the apex (models 3 or 4), rather than due to differences in *FT* signal generation in the leaf, and that this difference is due to the precocious endogenous developmental age pathway in the apex.

## Discussion

Research into the mechanisms of regulation of the floral transition has focussed largely on the model organism Arabidopsis. This has generated a demand for methods for translating this knowledge to other species. Here we demonstrate that apparently large differences in gene expression profiles over development between the closely related crop *B. rapa* and Arabidopsis can mostly be resolved through the application of a curve registration step during data analysis. We found that different genes require different registration functions, consistent with the desynchronisation of multiple regulatory modules within the GRN between these species. We identified exemplar differences in the regulation of the floral integrator gene *SOC1* between rapid cycling Arabidopsis and *B. rapa* in these developmental time-courses. Through comparison of gene expression profiles in R-o-18 and Sarisha-14, we have identified a putative *FT*-independent mechanism which potentiates the extremely early floral transition in Sarisha-14 and consequently underlies its commercial viability in Bangladesh.

### Flowering GRN in Col-0 vs R-o-18

Comparison of optimal registration functions for expression profiles of key floral genes indicates that expression of *SOC1* is delayed in *B. rapa* vs Arabidopsis relative to other gene expression profiles under these environmental conditions. Detailed comparison of patterns between gene expression profiles using the CSI algorithm identified differences in the relationships between expression of *FLC* and *FUL* and *SOC1*. In Arabidopsis, *SOC1* is partly regulated by the balance of FLC and FUL which compete to dimerise with SVP. FLC-SVP represses *SOC1* expression, whereas the FUL-SVP dimer activates it (Balanzà *et al*., 2014). Over time, apical *FLC* expression declines and *FUL* expression increases to the point that FUL-SVP becomes the dominant dimer. Gene regulatory links inferred from Arabidopsis gene expression data are consistent with this model, however those from *B. rapa* are not. Instead, in *B. rapa*, Bra*FLC* expression remains high until after *BraSOC1* expression is well established, even though it appears to encode a functional protein.

R-o-18 is commonly used as a model Brassica accession due to its rapid lifecycle and lack of vernalisation requirement, yet this analysis suggests that it could potentially be made to flower more rapidly. An interesting breeding objective to achieve this end would be to knock out expression of the *BraFLC*.*A03a* copy in the apex. We hypothesis that this may reduce competition for SVP dimerization, and allow precocious upregulation of *SOC1* expression, and subsequent changes in the regulation of its downstream target genes.

### Flowering time in Sarisha-14 vs Ro18

In Arabidopsis, flowering can be triggered under long-day, inductive conditions by FT, or by aging and phytohormones under non-inductive, short-day conditions (Hyun *et al*., 2016, 2019). Differences in the timing of the floral transition between R-o-18 and Sarisha-14 are not accounted for by differences in the expression profiles of *FT* homologues, which have similar expression patterns and levels until much later in development. Many components of the aging pathway to floral transition are under post-transcriptional control **(Fig. 6)**, and so not directly identifiable by RNA-seq. It is striking that gene expression of those key components which can be detected are consistent with differences in this pathway between Sarisha-14 and R-o-18.

Previous studies have identified a transposon insertion in the second intron of R-o-18 *BraFT*.*A7* which causes a reduction of expression as underlying a QTL between R-o-18 and the fast flowering caixin type L58 (Zhang *et al*., 2015). However, whilst in L58, similar expression levels were observed for both copies of *BraFT*, in Sarisha-14 we observed the same reduced expression of *BraFT*.*A7* as in R-o-18, indicating that this allele does not underlie the difference between Sarisha-14 and R-o-18.

*FT* expression is known to vary over the course of a day in Arabidopsis (Krzymuski *et al*., 2015; Song *et al*., 2018). Although samples from both varieties were taken at the same time, it is possible that differences in the expression dynamics over the diurnal cycle contribute to differences in development. It is also possible that potential differences in FT signalling effectiveness, or in tissue conductivity to long distance signals, contribute to differences in FT activity at the apex, which cannot be seen in gene expression level in the leaf. However, as we see no evidence for differences in the *FT* coding sequence between Sarisha-14 and R-o-18, and we do see evidence for differences in phytohormone and age-related signalling. Consequently, differences in the GRN at the apex is the most parsimonious explanation for the early flowering phenotype.

Interestingly, selective breeding appears to have produced a variety, Sarisha-14, that uses the aging GRN to trigger early flowering. The aging GRN can be viewed as an endogenous timer that normally acts in older meristems to allow flowering in the absence of FT under unfavourable environmental conditions (Hyun *et al*., 2019). In Sarisha-14 however, it apparently proceeds so rapidly that it becomes a trigger for flowering even under inductive, long-day environmental conditions, either in the absence of *FT*, or under lower concentration than is required in R-o-18. A challenge in determining the causal genomic differences between R-o-18 and Sarisha-14 is that the identified GRN is highly connected, incorporating post-transcriptional regulation and many key developmental phytohormones and sugar signalling into the regulation of aging. Identifying the causal alleles will, therefore, likely require use of a recombinant inbred line population.

## Conclusions

Flowering time control is of major importance in crop adaptation to different environments. Our study provides gene expression data for all genes in leaf and apex for two rapid cycling oil type *B. rapa* lines through the floral transition. By curve registration of gene expression profiles, and network inference, we have identified differences in the regulation of the floral transition between Arabidopsis and *B. rapa*. We also identified regulatory differences between *B. rapa* varieties and linked these to phenotypic differences. This demonstrates that GRNs differ even between closely related cultivars. The data presented provide a foundation for future breeding efforts of *B. rapa* crops.

## Availability of supporting data

The Illumina sequence reads have been deposited in NCBI Sequence Read Archive, project ID PRJNA593493.

## Acknowledgements

We are grateful to Dr Lei Zhang for providing an updated identification of paralogous *B. rapa* genes and mapping to Arabidopsis homologues. We thank Profs Lars Ostergaard and Steve Penfield for critical reading, valuable comments and suggestions. The authors acknowledge financial support from the Global Challenges Research Fund grant “Optimising maturity for enhanced yield in the Sarisha crop, Bangladesh” (BB/P01531X/1), and Biotechnology and Biological Sciences Research Council grants; Brassica rapeseed and vegetable optimisation strategic Lola (BB/P003095/1), and “Genes in the environment” (BB/P013511/1).

## Author Contributions

A.C. designed and performed the majority of the data analysis. D.M.J. provided technical support. J.H., E.T., S.W., L.B., A.C., J.I. & R.W. performed the experiments. M.A. & C.D. provided experimental material. R.J.M., R.W. & J.I. supervised the project. A.C. planned and wrote the first draft of the manuscript. A.C., R.W., J.I., J.H. and R.J.M. wrote the manuscript with contributions from all authors. All authors provided critical feedback that helped shape the analysis and the manuscript.

## Supporting information

**Table S1:** Query FASTA sequences, and BLAST hit locations in the Chiifu v3 reference sequence for precursor microRNA sequences.

**Table S2:** Gene ontology enrichment among groups of *B. rapa* genes which are best aligned to the Arabidopsis gene expression profile through different registration shift functions.

**Fig. S1:** The CSI network inference algorithm performs well on synthetic data similar to the experimental gene expression time-course.

**Fig. S2:** The number of genes identified as having similar gene expression in both organisms is significant in the real data.

**Fig. S3:** The distribution of identified optimal registration function parameters

**Fig. S4:** Differences in the CSI inferred regulation of *SOC1* between Arabidopsis and R-o-18.

**Fig. S5:** Gene expression profiles of *FLC* paralogues in R-o-18, expression of *BraFLC*.*A3a* (BRAA03G004170.3C) is dominant to the other *FLC* copies.

**Fig. S6:** Gene expression profiles of *FUL, FLC*, and *SOC1* in Arabidopsis and R-o-18.

**Fig. S7:** Apices of Sarisha-14 have floral morphology by day 10, R-o-18 has floral morphology by day 17.

**Fig. S8** CSI inferred evidence for regulatory relationships between genes expressed in the leaf, and floral integrator genes *SOC1* and *FUL*

**File S1:** *B. rapa* gene models identified by Stringtie v1.2.2 using time-course gene expression data, .gtf file format.

## References

Abe M, Kobayashi Y, Yamamoto S, Daimon Y, Yamaguchi A, Ikeda Y, Ichinoki H, Notaguchi M, Goto K, Araki T. 2005. FD, a bZIP protein mediating signals from the floral pathway integrator FT at the shoot apex. Science (New York, N.Y.) 309: 1052–6.

Achard P, Cheng H, De Grauwe L, Decat J, Schoutteten H, Moritz T, Van Der Straeten D, Peng J, Harberd NP. (2006). Integration of plant responses to environmentally activated phytohormonal signals. Science (New York, N.Y.) 311: 91–4.

Andrés F, Coupland G. 2012. The genetic basis of flowering responses to seasonal cues. Nature Reviews Genetics 13: 627–639.

Aukerman MJ, Sakai H. 2003. Regulation of flowering time and floral organ identity by a MicroRNA and its APETALA2-like target genes. The Plant Cell 15: 2730–41.

Balanzà V, Martínez-Fernández I, Ferrándiz C. 2014. Sequential action of FRUITFULL as a modulator of the activity of the floral regulators SVP and SOC1. Journal of Experimental Botany 65: 1193–203.

Beilstein MA, Nagalingum NS, Clements MD, Manchester SR, Mathews S. 2010. Dated molecular phylogenies indicate a Miocene origin for Arabidopsis thaliana. Proceedings of the National Academy of Sciences of the United States of America 107: 18724–18728.

Berardini TZ, Reiser L, Li D, Mezheritsky Y, Muller R, Strait E, Huala E. 2015. The arabidopsis information resource: Making and mining the ‘gold standard’ annotated reference plant genome. Genesis 53: 474–485.

Bernier G, Périlleux C. 2005. A physiological overview of the genetics of flowering time control. Plant Biotechnology Journal 3: 3–16.

Blümel M, Dally N, Jung C. 2015. Flowering time regulation in crops — what did we learn from Arabidopsis? Current Opinion in Biotechnology 32: 121–129.

Bouché F, D’Aloia M, Tocquin P, Lobet G, Detry N, Périlleux C. 2016a. Integrating roots into a whole plant network of flowering time genes in Arabidopsis thaliana. Scientific Reports 6: 29042.

Bouché F, Lobet G, Tocquin P, Périlleux C. 2016b. FLOR-ID: an interactive database of flowering-time gene networks in *Arabidopsis thaliana*. Nucleic Acids Research 44: D1167–D1171.

Canola Council of Canada. 2013. Canola Encyclopedia: time-of-seeding.

Chen X. 2004. A MicroRNA as a Translational Repressor of APETALA2 in Arabidopsis Flower Development. Science 303: 2022–2025.

Conti L. 2017. Hormonal control of the floral transition: Can one catch them all? Developmental Biology 430: 288–301.

Corbesier L, Vincent C, Jang S, Fornara F, Fan Q, Searle I, Giakountis A, Farrona S, Gissot L, Turnbull C, et al. 2007. FT Protein Movement Contributes to Long-Distance Signaling in Floral Induction of Arabidopsis. Science 316: 1030–1033.

Franks SJ, Perez-Sweeney B, Strahl M, Nowogrodzki A, Weber JJ, Lalchan R, Jordan KP, Litt A. 2015. Variation in the flowering time orthologs BrFLC and BrSOC1 in a natural population of Brassica rapa. PeerJ 3: e1339.

Fu X, Harberd NP. 2003. Auxin promotes Arabidopsis root growth by modulating gibberellin response. Nature 421: 740–743.

Griffiths-Jones S, Grocock RJ, van Dongen S, Bateman A, Enright AJ. 2006. miRBase: microRNA sequences, targets and gene nomenclature. Nucleic Acids Research 34: D140–D144.

He J, Xu M, Willmann MR, McCormick K, Hu T, Yang L, Starker CG, Voytas DF, Meyers BC, Poethig RS. 2018. Threshold-dependent repression of SPL gene expression by miR156/miR157 controls vegetative phase change in Arabidopsis thaliana. PLOS Genetics 14: e1007337.

Hong Y, Jackson S. 2015. Floral induction and flower formation-the role and potential applications of miRNAs. Plant Biotechnology Journal 13: 282–292.

Hyun Y, Richter R, Vincent C, Martinez-Gallegos R, Porri A, Coupland G. 2016. Multi-layered Regulation of SPL15 and Cooperation with SOC1 Integrate Endogenous Flowering Pathways at the Arabidopsis Shoot Meristem. Developmental Cell 37: 254–266.

Hyun Y, Vincent C, Tilmes V, Bergonzi S, Kiefer C, Richter R, Martinez-Gallegos R, Severing E, Coupland G. 2019. A regulatory circuit conferring varied flowering response to cold in annual and perennial plants. Science (New York, N.Y.) 363: 409–412.

Ims RA. 1990. The ecology and evolution of reproductive synchrony. Trends in Ecology & Evolution 5: 135–140.

Jaeger KE, Wigge PA. 2007. FT Protein Acts as a Long-Range Signal in Arabidopsis. Current Biology 17: 1050–1054.

Kim D, Langmead B, Salzberg SL. 2015. HISAT: a fast spliced aligner with low memory requirements. Nature Methods 12: 357–60.

Klepikova A V., Logacheva MD, Dmitriev SE, Penin AA. 2015. RNA-seq analysis of an apical meristem time series reveals a critical point in Arabidopsis thaliana flower initiation. BMC Genomics 16: 466.

Krijthe JH. 2015. Rtsne: T-Distributed Stochastic Neighbor Embedding using Barnes-Hut Implementation.

Krzymuski M, Andres F, Cagnola JI, Jang S, Yanovsky MJ, Coupland G, Casal JJ. 2015. The dynamics of FLOWERING LOCUS T expression encodes long-day information. The Plant Journal 83: 952–961.

Lee J, Lee I. 2010. Regulation and function of SOC1, a flowering pathway integrator. Journal of Experimental Botany 61: 2247–2254.

Lee T, Yang S, Kim E, Ko Y, Hwang S, Shin J, Shim JE, Shim H, Kim H, Kim C, et al. 2015. AraNet v2: an improved database of co-functional gene networks for the study of Arabidopsis thaliana and 27 other nonmodel plant species. Nucleic Acids Research 43: D996–D1002.

Leiboff S, Hake S. 2019. Reconstructing the Transcriptional Ontogeny of Maize and Sorghum Supports an Inverse Hourglass Model of Inflorescence Development. Current Biology 29: 3410–3419.e3.

Lorrai R, Boccaccini A, Ruta V, Possenti M, Costantino P, Vittorioso P. 2018. Abscisic acid inhibits hypocotyl elongation acting on gibberellins, DELLA proteins and auxin. AoB Plants 10.

Lou P, Zhao J, Kim JS, Shen S, Del Carpio DP, Song X, Jin M, Vreugdenhil D, Wang X, Koornneef M, et al. 2007. Quantitative trait loci for flowering time and morphological traits in multiple populations of Brassica rapa. Journal of Experimental Botany 58: 4005–4016.

van der Maaten LJP, Hinton GE. 2008. Visualizing High-Dimensional Data Using t-SNE. Journal of Machine Learning Research 9: 2579–2605.

McClung CR, Lou P, Hermand V, Kim JA. 2016. The Importance of Ambient Temperature to Growth and the Induction of Flowering. Frontiers in Plant Science 7: 1–7.

Md MUH, Md NI, Kadir M, Miah NH. 2016. Performance of rapeseed and mustard (Brassica sp.) varieties/lines in north-east region (Sylhet) of Bangladesh. Advances in Plants & Agriculture Research 5.

Meier U, Bleiholder H, Buhr L, Feller C, Hack H, Heß M, Lancashire P., Schnock U, Stauß R, Van Den Boom T, et al. 2009. The BBCH system to coding the phenological growth stages of plants–history and publications. Journal für Kulturpflanzen 61: 41–52.

Mia AB. 2017. Digital herbarium of crop plants.

Miah MAM, Mondal MRI. 2017. Oilseeds sector of Banglasesh: challenges and opportunities. SAARC Journal of Agriculture 15: 161–172.

Newman M. 2010. Networks : an introduction. New York, New York, USA: Oxford University Press.

del Olmo I, Poza-Viejo L, Piñeiro M, Jarillo JA, Crevillén P. 2019. High ambient temperature leads to reduced FT expression and delayed flowering in Brassica rapa via a mechanism associated with H2A.Z dynamics. The Plant Journal 100: 343–356.

Pajoro A, Biewers S, Dougali E, Leal Valentim F, Mendes MA, Porri A, Coupland G, Van De Peer Y, Van Dijk ADJ, Colombo L, et al. 2014. The (r)evolution of gene regulatory networks controlling Arabidopsis plant reproduction; a two decades history. Journal of Experimental Botany 65: 4731–4745.

Penfold CA, Wild DL. 2011. How to infer gene networks from expression profiles, revisited. Interface Focus 1: 857–870.

Périlleux C, Bouché F, Randoux M, Orman-Ligeza B. 2019. Turning Meristems into Fortresses. Trends in Plant Science 24: 431–442.

Pertea M, Pertea GM, Antonescu CM, Chang T-C, Mendell JT, Salzberg SL. 2015. StringTie enables improved reconstruction of a transcriptome from RNA-seq reads. Nature Biotechnology 33: 290–295.

Ramsay J, Silverman BW. 2005. The registration and display of functional data. In: Functional Data Analysis. New York, New York, USA: Springer, 127–146.

Rana D, Boogaart T, O’Neill CM, Hynes L, Bent E, Macpherson L, Park JY, Lim YP, Bancroft I. 2004. Conservation of the microstructure of genome segments in Brassica napus and its diploid relatives. The Plant Journal 40: 725–733.

Robinson MD, McCarthy DJ, Smyth GK. 2010. edgeR: a Bioconductor package for differential expression analysis of digital gene expression data. Bioinformatics 26: 139–140.

Schiessl S V., Huettel B, Kuehn D, Reinhardt R, Snowdon RJ. 2017. Flowering Time Gene Variation in Brassica Species Shows Evolutionary Principles. Frontiers in Plant Science 8: 1–13.

Simpson GG, Dean C. 2002. Arabidopsis, the Rosetta stone of flowering time? Science 296: 285–289.

Song YH, Kubota A, Kwon MS, Covington MF, Lee N, Taagen ER, Laboy Cintrón D, Hwang DY, Akiyama R, Hodge SK, et al. 2018. Molecular basis of flowering under natural long-day conditions in Arabidopsis. Nature Plants 4: 824–835.

Wu G, Poethig RS. 2006. Temporal regulation of shoot development in Arabidopsis thaliana by miR156 and its target SPL3. Development 133: 3539–3547.

Wu J, Wei K, Cheng F, Li S, Wang Q, Zhao J, Bonnema G, Wang X. 2012. A naturally occurring InDel variation in BraA.FLC.b (BrFLC2) associated with flowering time variation in Brassica rapa. BMC Plant Biology 12: 1–9.

Xiao D, Zhao JJ, Hou XL, Basnet RK, Carpio DPD, Zhang NW, Bucher J, Lin K, Cheng F, Wang XW, et al. 2013. The Brassica rapa FLC homologue FLC2 is a key regulator of flowering time, identified through transcriptional co-expression networks. Journal of Experimental Botany 64: 4503–4516.

Yant L, Mathieu J, Dinh TT, Ott F, Lanz C, Wollmann H, Chen X, Schmid M. 2010. Orchestration of the floral transition and floral development in Arabidopsis by the bifunctional transcription factor APETALA2. The Plant Cell 22: 2156–70.

Yoo SK, Chung KS, Kim J, Lee JH, Hong SM, Yoo SJ, Yoo SY, Lee JS, Ahn JH. 2005. CONSTANS activates SUPPRESSOR OF OVEREXPRESSION OF CONSTANS 1 through FLOWERING LOCUS T to promote flowering in Arabidopsis. Plant Physiology 139: 770–778.

Yu S, Galvão VC, Zhang Y-C, Horrer D, Zhang T-Q, Hao Y-H, Feng Y-Q, Wang S, Schmid M, Wang J-W. 2012. Gibberellin Regulates the Arabidopsis Floral Transition through miR156-Targeted SQUAMOSA PROMOTER BINDING–LIKE Transcription Factors. The Plant Cell 24: 3320–3332.

Yuan Y-X, Wu J, Sun R-F, Zhang X-W, Xu D-H, Bonnema G, Wang X-W. 2009. A naturally occurring splicing site mutation in the Brassica rapa FLC1 gene is associated with variation in flowering time. Journal of Experimental Botany 60: 1299–1308.

Zhang L, Cai X, Wu J, Liu M, Grob S, Cheng F, Liang J, Cai C, Liu Z, Liu B, et al. 2018. Improved Brassica rapa reference genome by single-molecule sequencing and chromosome conformation capture technologies. Horticulture Research 5: 1–11.

Zhang X, Meng L, Liu B, Hu Y, Cheng F, Liang J, Aarts MGM, Wang X, Wu J. 2015. A transposon insertion in FLOWERING LOCUS T is associated with delayed flowering in Brassica rapa. Plant Science 241: 211–220.

