## Supplemental Figure S1 for "Comparative transcriptomics identifies differences in the regulation of the floral transition between Arabidopsis and *Brassica rapa* cultivars"

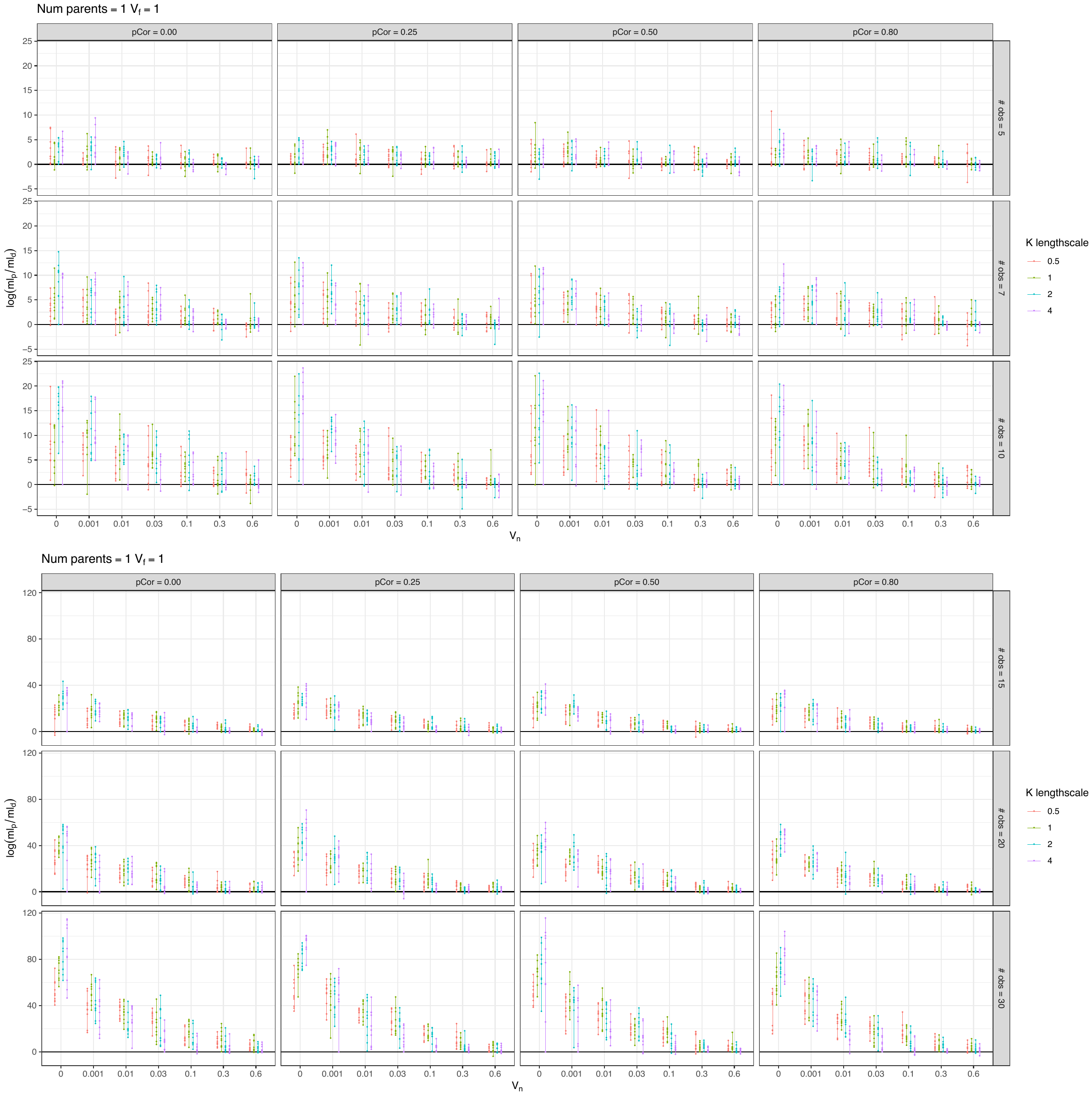

**Fig. S1:**

**The CSI network inference algorithm performs well on synthetic data similar to the experimental gene expression time-course.**

Synthetic gene expression data was generated for a “true parent”, a “dummy parent”, and a “child” gene which is regulated by the “true parent”. “true parent”, and “dummy parent” gene expression data were sampled from a multivariate gaussian distribution. “Child” gene expression data was sampled from a Gaussian Process, over “true parent” gene expression (with function variance, V_f_ =1 and length-scale=K). Gaussian noise was added to child gene expression, sampled from N(0, V_n_).

The marginal likelihood for regulation of the child gene by the “true parent” (ml_p_), and “dummy parent” (ml_d_) were inferred using the CSI algorithm. Each point is a result using separate synthetic dataset. Vertical bars join best and worst performing run for given generative parameters. pCor is the correlation between “true parent” and “dummy parent” expression. “# obs” is the number of observations of gene expression for “parent”, “dummy”, and “child” genes.

Simulated results suggest that the CSI algorithm is well able to identify the true, generative parent given 30 observations, even with large amounts of added noise (V_n_). This number of observations is similar to our experimental data (31 observations of *A. thaliana* Col-0, 33 observations of *B. rapa* R-o-18). The true noise and length-scale is difficult to determine in the real experimental data, but CSI performs well over a broad range of values.
