## Supplemental Figure S2 for "Comparative transcriptomics identifies differences in the regulation of the floral transition between Arabidopsis and *Brassica rapa* cultivars"

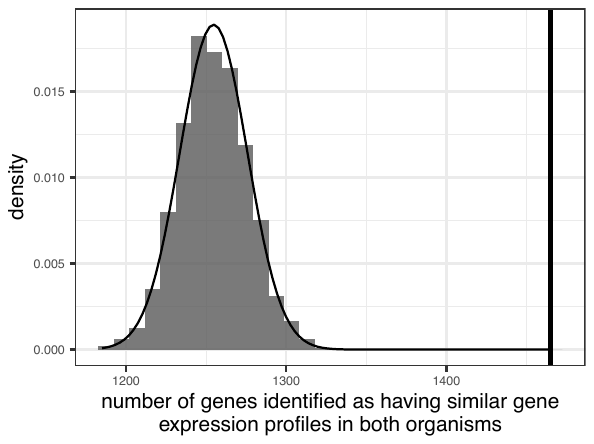

**Fig S2**:

**The number of genes identified as having similar gene expression in both organisms is significant in the real data.** Genes with similar gene expression in both organisms were identified as genes in which a registration model better explains the gene expression data than separate models, as described in Methods. Here, the number of genes identified in the real data (vertical line) are compared to the number of genes identified in 1000 random permutations of the data in which each gene’s expression in one organism is compared to a randomly allocated gene in the other. None of the permuted trials were found to have an equal or greater number of genes for which the registration model was preferred, indicating significance at the p < 0.001 level, though it can be seen from the distribution of results, that this value is likely far smaller. Assuming a normal distribution for the permuted results (plotted) suggests a p value < 1e-23. Interestingly, a large number of genes are identified, even when comparisons are made between random genes. Anecdotally, this appears to be at least partly because many genes share similar gene expression profiles, resulting in somewhat frequent comparison to a random gene with a similar expression profile. It is additionally presumed that there is some overfitting during the registration process, though the balance between these two causes is difficult to determine.
