## Supplemental Figure S3 for "Comparative transcriptomics identifies differences in the regulation of the floral transition between Arabidopsis and *Brassica rapa* cultivars"

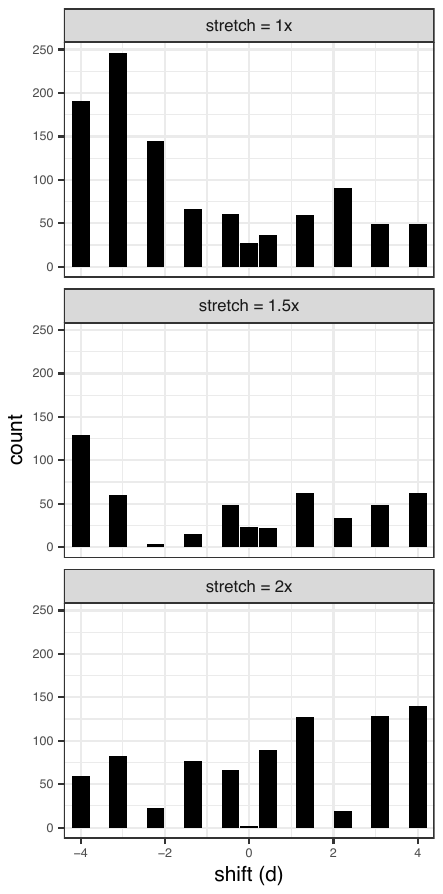

**Fig. S3:**

**Distribution of identified optimal registration function parameters.** A wide variety of different optimal parameters are identified for different genes, indicating that many differently synchronised processed occur, though some appear more common than others.
