## Supplemental Figure S4 for "Comparative transcriptomics identifies differences in the regulation of the floral transition between Arabidopsis and *Brassica rapa* cultivars"

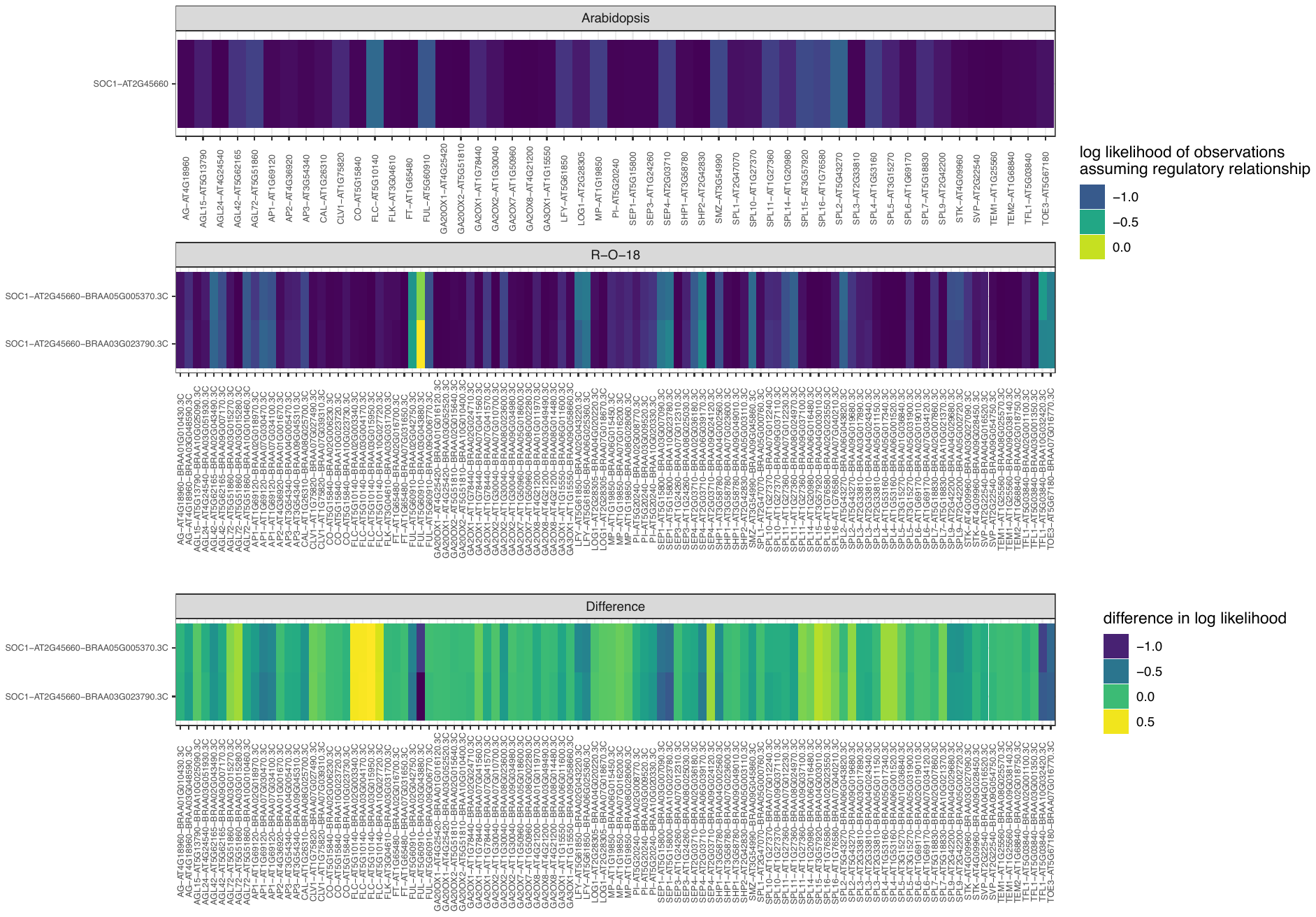

**Fig. S4:**

**Large differences in the gene expression evidence for regulation of *SOC1* by *FLC* and *FUL* exist between Arabidopsis and R-o-18**.

The likelihood of regulation of *SOC1* by candidate regulatory genes were calculated based on gene expression profiles via the Causal Structure Inference (CSI) algorithm as described in methods. Plot shows CSI calculated likelihood of observed data, assuming a regulatory link between *SOC1* and the candidate regulator genes in **a)** Col-0, **b)** R-o-18, & **c)** the difference between them. This is considered proportional to the likelihood of a regulatory link given the gene expression data. Among the candidate regulators of *SOC1*, some of the largest differences between species in the magnitude of the likelihood were for *FLC*, and *FUL* *A02* and *A03* gene copies.
