## Supplemental Figure S5 for "Comparative transcriptomics identifies differences in the regulation of the floral transition between Arabidopsis and *Brassica rapa* cultivars"

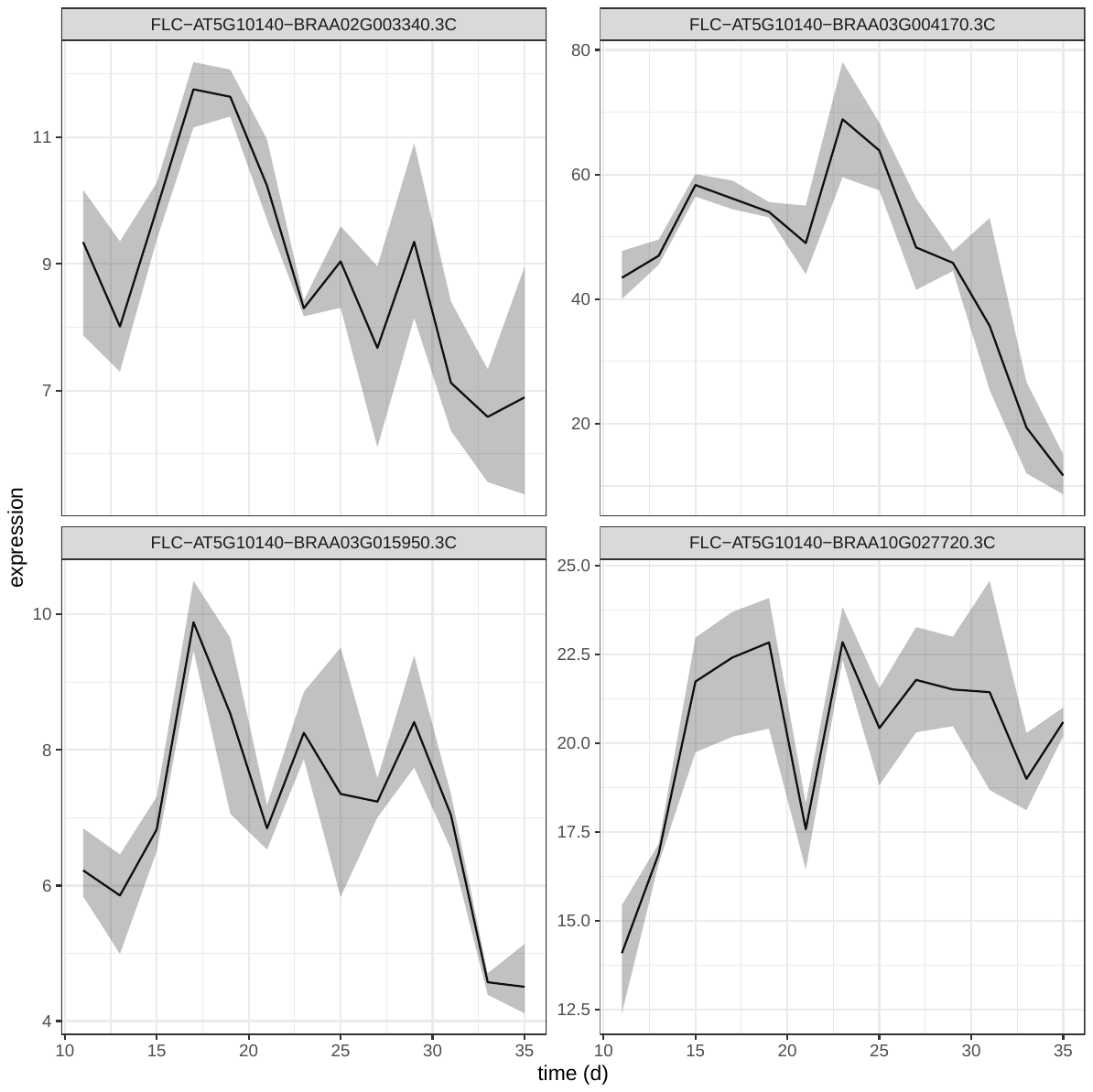

**Fig. S5:**

**Gene expression profiles of *FLC* paralogues in R-o-18,** expression of *FLC.A3a* (BRAA03G004170.3C) is dominant to the other FLC paralogues.
