## Supplemental Figure S6 for "Comparative transcriptomics identifies differences in the regulation of the floral transition between Arabidopsis and *Brassica rapa* cultivars"

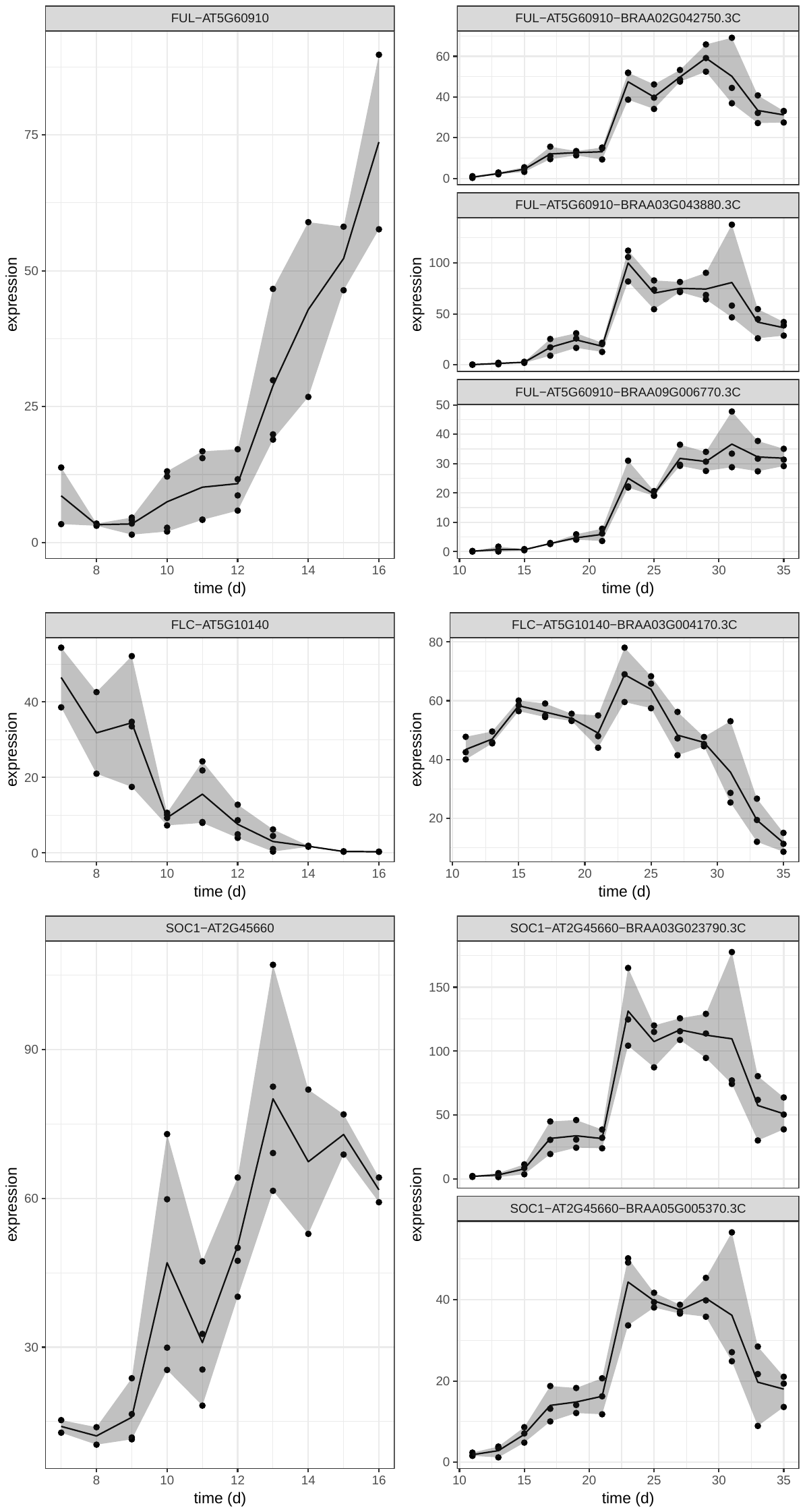

**Fig. S6:**

**Gene expression profiles of *FUL*, *FLC*, and *SOC1* in Arabidopsis and R-o-18**. Unlike in R-o-18, *FLC* expression declines before floral transition in Arabidopsis. In the Arabidopsis dataset, floral transition was determined to occur approximately 10d after germination (Klepikova *et al.*, 2015). In R-o-18, transition occurred by 17d. Expression is EdgeR normalised cpm. Brassica orthologues of Arabidopsis genes identified based on annotation in Chiifu v3 reference sequence (Zhang *et al.*, 2018).
