## Supplemental Figure S7 for "Comparative transcriptomics identifies differences in the regulation of the floral transition between Arabidopsis and *Brassica rapa* cultivars"

### Slide 1
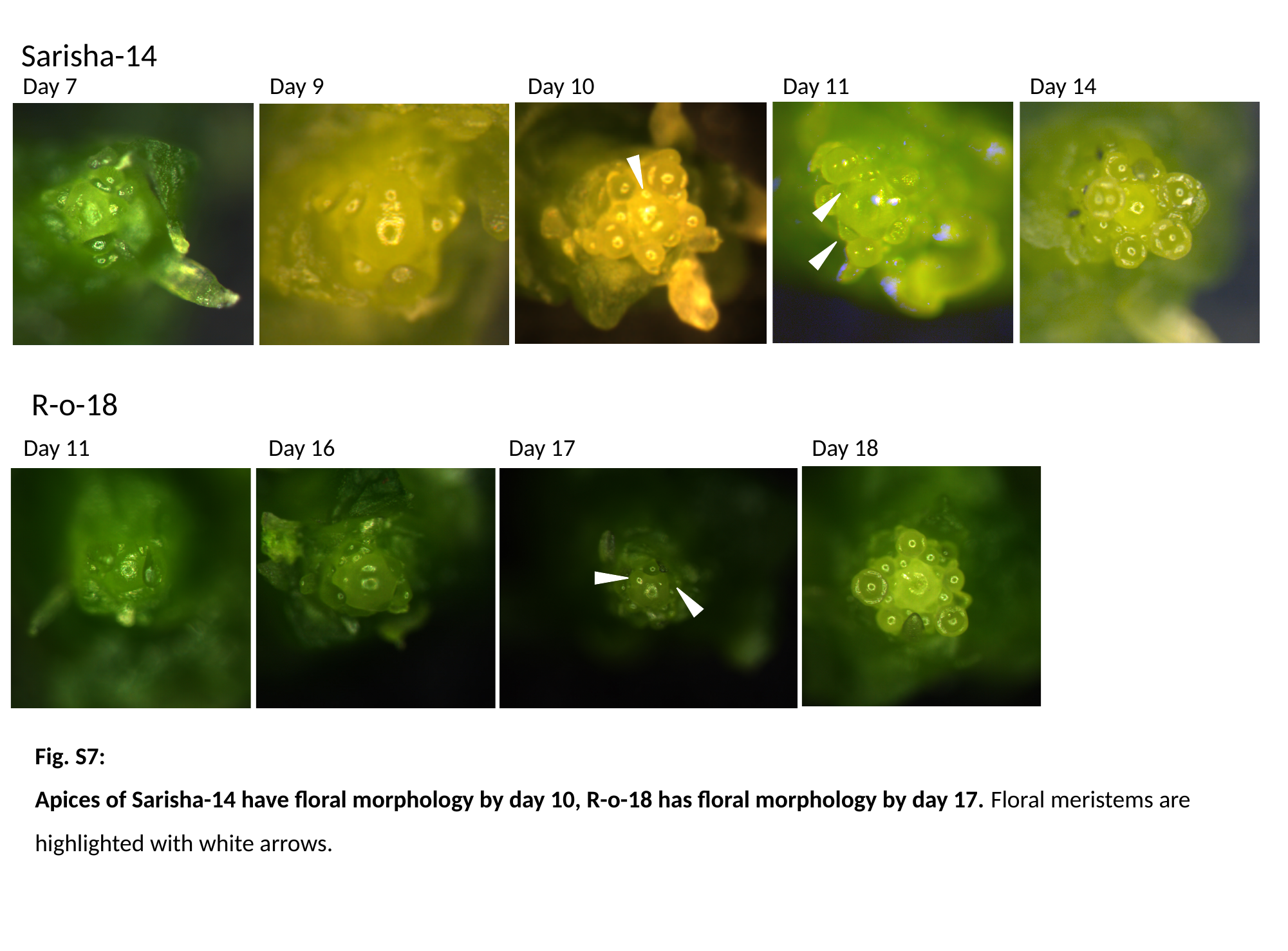

Sarisha-14
Day 7
Day 9
Day 10
Day 11
Day 14
R-o-18
Day 11
Day 16
Day 17
Day 18
Fig. S7:
Apices of Sarisha-14 have floral morphology by day 10, R-o-18 has floral morphology by day 17. Floral meristems are highlighted with white arrows.
