## Supplemental Figure S8 for "Comparative transcriptomics identifies differences in the regulation of the floral transition between Arabidopsis and *Brassica rapa* cultivars"

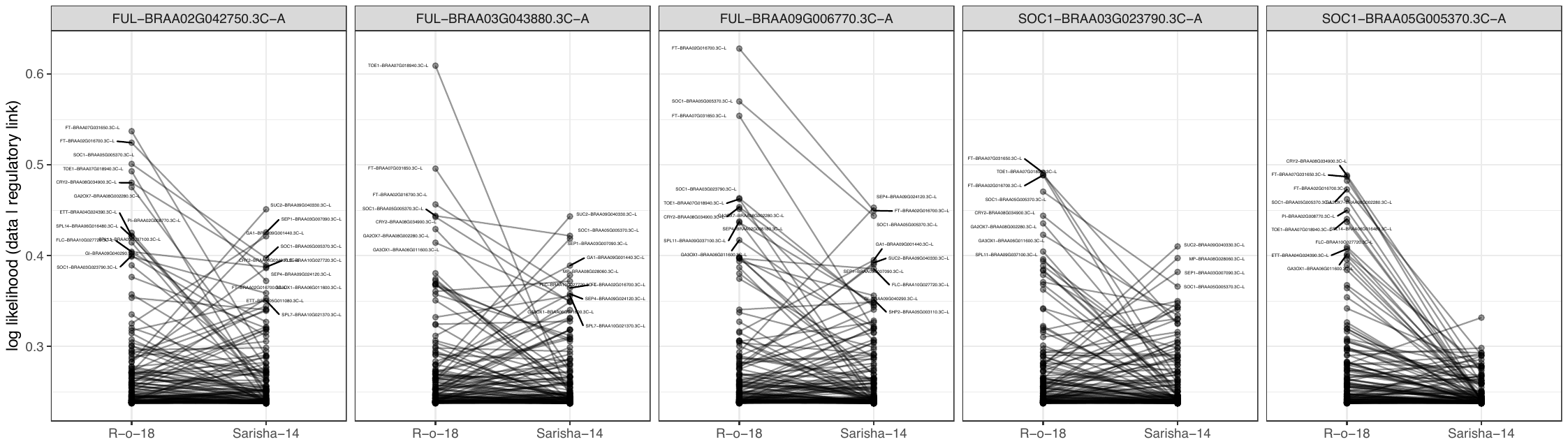

**Fig S8: CSI inferred evidence for regulatory relationships between genes expressed in the leaf, and floral integrator genes *SOC1* and *FUL* expressed in the apex*.*** The likelihood of observing the gene expression data, given a regulatory link between gene expression profiles in the leaf of R-o-18 and Sarisha-14, and the expression of *SOC1*, and *FUL* in the apex is calculated using the CSI network inference algorithm. This is proportional to the probability of a regulatory link given no prior information. Stronger evidence for regulatory links between the leaf and apex are found in R-o-18 than in Sarisha-14. In particular, expression of *FT* in the leaf is more associated with expression of *SOC1* and *FUL* in the apex. Other genes, for example the sugar transported *SUC2* are more associated with gene expression in the apex than *FT* in Sarisha-14.
